# Individual-Specific Gaussian Graphical Models for Heterogeneous Populations with Application to Epigenetic Gene Regulation in Lung Adenocarcinoma

**DOI:** 10.64898/2026.05.25.727641

**Authors:** Enakshi Saha

## Abstract

Inter-patient molecular heterogeneity is a fundamental challenge in precision oncology: population-level multi-omics networks reveal average biology aggregated across the population but obscure individual variations that drive differential clinical outcomes. We introduce SIREN (Sample-specific Inference via Regularized Empirical-Bayes Networks), a method that estimates one partial correlation network per sample across omics layers by combining a population-level empirical Bayes prior with a rank-1 individual-specific update. Since a sample-specific precision matrix cannot be estimated from a single observation, SIREN uses a conjugate Inverse Wishart prior whose mean is the Oracle Approximating Shrinkage estimator, yielding closed-form individual-specific posteriors without MCMC. On simulated heterogeneous populations, SIREN achieves superior edge recovery over population-average methods including OAS, Ledoit-Wolf, and graphical Lasso, while remaining competitive in homogeneous settings. Applied to paired transcriptomic and methylomic profiles from lung adenocarcinoma, SIREN identifies individual-specific gene-methylation regulatory edges that stratify patients by survival in ways population-level analysis cannot, implicating chromatin remodeling and WNT signaling pathways in epigenetic heterogeneity. SIREN is computationally scalable and available as a Python package.

## 1 Introduction

Lung adenocarcinoma (LUAD) is the most common subtype of non-small cell lung cancer and one of the leading causes of cancer-related mortality worldwide (Kanchustambham & Sharma 2026). Despite advances in staging, and treatment, patients diagnosed at the same clinical stage often exhibit markedly different survival outcomes, suggesting that individual-level biological factors beyond tumor stage, which is typically determined by the size and spread of tumor, drive prognosis. A growing body of evidence implicates epigenetic dysregulation, particularly aberrant DNA methylation as a key driver of inter-patient heterogeneity in cancer (Gimeno-Valiente et al. 2025). Understanding how DNA methylation-driven gene regulatory relationships vary across individuals, and how these individual differences relate to survival, could reveal new therapeutic targets and guide patient stratification for epigenetic therapies (Bates 2020).

Gaussian Graphical Models (GGMs) have been extensively used for estimating gene regulatory networks from omics data, where specific edges represent partial correlations between molecular variables after controlling for all other variables in the network (Tian et al. 2016, Kotiang & Eslami 2020). However, these methods estimate a single network shared across all individuals, obscuring the individual-level heterogeneity in how different molecular variables interact with each other, a phenomenon critical to understanding why individuals with the same disease diagnosis have varying prognosis and response to therapy. To our knowledge, no existing method estimates individual-specific Gaussian graphical models from omics data, a critical gap given the well-documented biological heterogeneity across cancer patients.

Several methods have been proposed for estimating Gaussian graphical models in heterogeneous populations. Joint graphical lasso approaches (Danaher et al. 2014, Guo et al. 2011) estimate multiple precision matrices simultaneously for observations belonging to multiple groups but these methods require the known group membership of each observation. Gao et al. (2016) and Hao et al. (2018) relax this assumption by simultaneously clustering samples and estimating group-level precision matrices, while Wu et al. (2024) proposed a heterogeneous latent transfer learning approach that identifies latent subpopulation structures across multiple studies and transfers knowledge within the same subpopulation. Despite these advances, all aforementioned methods estimate one precision matrix per subpopulation rather than per individual, making no distinction between patients assigned to the same subgroup, which is a critical limitation in precision medicine applications where each patient may have a unique gene regulatory network driving heterogeneity in disease mechanisms.

Individual-level gene regulatory networks have been used extensively in precision medicine applications in lung cancer (Saha et al. 2024a, Chen et al. 2025, Saha et al. 2025, Lopes-Ramos et al. 2026). These studies rely on individual-specific estimation of correlation matrices. However, these methods have several important caveats. For example, Yu et al. (2015), Kuijjer et al. (2019) and Chen et al. (2023) compute gene co-expression matrices that are not guaranteed to be positive semi-definite, often yielding negative eigenvalues and correlation values outside [−1, 1]. Recently, (Saha et al. 2024b) rectified these limitations by introducing BONOBO, a method for estimating individual-specific correlation networks using a leave-one-out sample covariance matrix as the prior for each individual and demonstrated the efficacy of individual-specific networks in identifying heterogeneous disease biology (Saha 2026). Building on BONOBO, Saha et al. (2026) proposed PRISM, a method for estimating co-expression matrices for cross-sectional and longitudinal omics data, providing superior coverage and computational efficiency. However, both BONOBO and PRISM produce correlation networks rather than partial correlation networks. Partial correlations, derived from the precision matrix (inverse covariance), capture direct regulatory relationships by controlling for all other variables, whereas correlations reflect both direct and indirect associations. Furthermore, individual-specific covariance matrices estimated by BONOBO are rank-deficient when *p* ≥ *n*, as the leave-one-out sample covariance has rank at most *n* − 1, and therefore cannot be inverted to obtain partial correlations.

To fill this gap, we propose SIREN (Sample-specific Inference via Regularized Empirical-Bayes Networks), an empirical Bayes framework for estimating individual-specific partial correlation networks from omics data. We assume each individual observations follows a Gaussian distribution with a distinct unknown covariance structure and impose an Inverse Wishart prior on each individual’s covariance matrix, with the prior mean estimated from all samples using the Oracle Approximating Shrinkage (OAS) estimator (Chen et al. 2010) following the empirical Bayes principle. The posterior mean of each individual’s precision matrix is then derived as a closed form weighted average of the prior mean and a rank-1 sample-specific update, yielding a computationally efficient estimator that shrinks individual covariance networks towards the population average, analogously to how James-Stein estimators shrink sample-specific maximum likelihood estimates of individual means towards the grand mean (Stein 1956, James et al. 1961).

We validate SIREN through two complementary analyses before applying it to our main lung cancer application. First, using simulation studies for varying dimensions and sample sizes, we demonstrate that SIREN outperforms population-level methods in heterogeneous populations while maintaining competitive performance in homogeneous settings. Second, using a yeast transcription factor (TF) knockout dataset (Jackson et al. 2020) we show that SIREN recovers known TF-target pathway biology, providing experimental ground truth validation of SIREN networks. Finally, we apply SIREN to paired gene expression and DNA methylation data from 448 patients with Lung Adenocarcinoma (LUAD) from The Cancer Genome Atlas (TCGA) cohort (Weinstein et al. 2013) and identify survival-associated epigenetic regulatory edges involving known cancer genes including *HMGA2, XRCC5, CKS2*, and *RUVBL2*. Through gene set enrichment analysis (GSEA) we show that epigenetic regulation of chromatin remodeling, WNT signaling, and stem cell differentiation pathways are associated with positive survival outcome, suggesting that individual differences in methylation-driven gene regulation may be a contributing factor to survival heterogeneity among lung cancer patients diagnosed at the same tumor stage.

The remainder of the paper is organized as follows. Section 2 describes the SIREN methodology. Section 3 presents applications on simulated datasets. Section 4 describes methodological validation on the yeast TF knockout experiment. Section 5 presents application on LUAD. Finally, Section 6 concludes with a discussion.

## 2 Methods

Gaussian Graphical Models (GGMs) represent conditional independence structure among a set of variables through an undirected graph, where the absence of an edge between two variables indicates conditional independence given all others. Formally, let **x** ∈ ℝ^*g*^ denote a *g*-dimensional random vector following a multivariate Gaussian distribution,

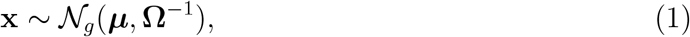

where ***µ*** ∈ ℝ^*g*^ is the mean vector and **Ω** ∈ ℝ^*g*×*g*^ is the precision matrix, i.e., the inverse of the covariance matrix **Σ** = **Ω**^−1^. The conditional independence structure of **x** is encoded directly in **Ω**: the (*k, l*)-th off-diagonal entry of **Ω** denoted by *ω*_*kl*_ is zero if and only if variables *k* and *l* are conditionally independent given all other remaining variables. The partial correlation between variables *k* and *l*, measuring their association after controlling for all others, is obtained by standardizing the off-diagonal entries of **Ω** as 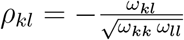. GGMs have therefore become a natural framework for estimating sparse dependency networks from high-dimensional data. However, in heterogeneous populations where individuals may have distinct dependency structures, a single population-level precision matrix estimate obscures this variation. Existing methods such as the graphical Lasso (Friedman et al. 2008), Ledoit-Wolf Estimator Ledoit & Wolf (2004) and Oracle Approximating Shrinkage (Chen et al. 2010) estimate a single **Ω** shared across all samples, collapsing individual-specific heterogeneity into a population average. Our goal is to estimate *individual-specific* precision matrices **Ω**_*i*_, one per observation *i* that capture this heterogeneity across individuals. The key challenge is that in most bulk-omics data each individual typically contributes only a single observation to the dataset, making such estimates statistically intractable under frequentist setup. We adopt an empirical Bayes approach motivated by the empirical Bayes formulation of the James-Stein type estimators (James et al. 1961, Efron & Morris 1972*a*).

### 2.1 Estimating Individual-specific Precision Matrix

Let us consider a set of *g* omics variables measured on *n* samples. Let 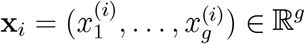 represent the *normalized* and *centered* observations for individual *i*; Let **X** represent the *n* × *g* matrix whose *i*^*th*^ row is **x**_*i*_. We assume that for every individual *i*, the centered omics vector follows a multivariate Normal distribution conditionally on its individual-specific covariance matrix:

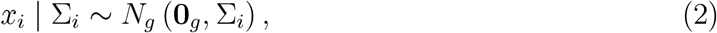

where **0**_*g*_ denotes a vector of all zeros Since each individual may have a distinct covariance structure, we treat Σ_*i*_ as a sample-specific unknown and place an Inverse Wishart prior on it:

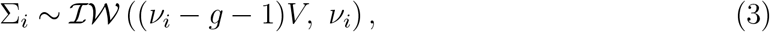

where *ν*_*i*_ ≥ *g* + 1 is the degrees of freedom parameter and *V* is a *g* × *g* positive definite matrix representing the prior mean of Σ_*i*_, since

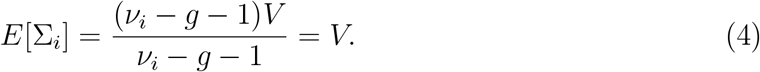

Marginalizing over Σ_*i*_, the implied marginal distribution of each centered observation is a multivariate *t*-distribution:

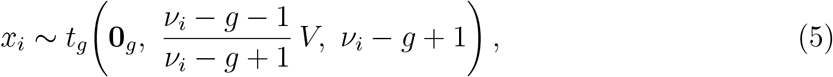

which reveals that *V* is the key population-level parameter governing the covariance structure shared across individuals and the sample covariance matrix 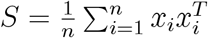 is an unbiased estimator of *V*. This provides a principled empirical Bayes justification for using *S* as the prior mean of *V*. However, in high-dimensional omics settings where *g* ≥ *n*, the sample covariance matrix *S* has rank at most *n* − 1 *< g* and is therefore singular, making it non-invertible. To overcome this, we follow Chen et al. (2010) and replace *S* with the shrinkage estimator:

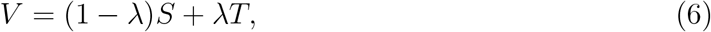

where 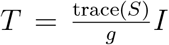 is a full-rank diagonal matrix whose diagonal entries are the mean variances across all omics variables, and *λ* ∈ (0, 1) is a shrinkage parameter. Since *T* is strictly positive definite, *V* is guaranteed to be invertible for any *λ* ∈ (0, 1).

Empirical Bayes estimators of the form (6) have been shown to dominate estimators of the form *aS* (Haff 1980, Ledoit & Wolf 2004, Chen et al. 2010), scalar multiples of the sample covariance *S*. As prior covariance, we choose the oracle approximating shrinkage (OAS) estimator proposed by (Chen et al. 2010). The OAS estimator provides a closed-form, data-driven estimate of the shrinkage parameter *λ* that minimizes the mean squared error of the covariance estimator under Gaussian assumptions, without requiring cross-validation or additional tuning. Moreover, Chen et al. (2010) demonstrate that the OAS estimator improves upon the widely used Ledoit–Wolf estimator (Ledoit & Wolf 2004), with particularly pronounced gains in the high-dimensional regime where *g* ≥ *n*.

The above model is equivalent to specifying that the precision matrix

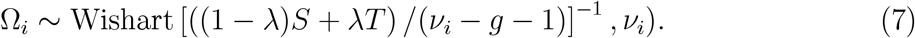

Due to the above assumption of multivariate normality, this model can only accommodate omics variables which can be transformed to approximately resemble a unimodal continuously distributed random variable. Several common molecular data types satisfy this requirement following appropriate preprocessing. Microarray-based gene expression data, after standard normalization and log2 transformation, exhibit approximately normal distributions. Similarly, RNA-sequencing count data, despite their discrete nature, follow approximate normality after normalization and log2 transformation (Saha et al. 2024b). DNA methylation M-values, defined as the log2 ratio of methylated to unmethylated signal intensities, likewise demonstrate approximately normal distributions.

#### Remark

For molecular modalities such as somatic mutations and copy number variations that cannot be easily transformed to approximate normality, a nonparanormal transformation (Liu et al. 2009) can be applied prior to model fitting, preserving the rank-based dependence structure among molecular variables.

Under this prior specification, the posterior distribution of the sample-specific precision matrix Ω_*i*_ also turns out to be Wishart, as described by the following theorem.

#### Theorem 2.1

*Under assumptions* (2) *and* (3), *the posterior distribution of* Σ_*i*_ *is*

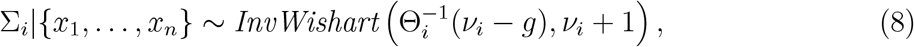

*where* 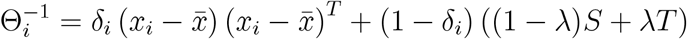 *denotes the posterior mean of* Σ_*i*_ *and* 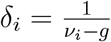.

Proof is provided in supplementary material S1.

The posterior mean of Σ_*i*_, denoted 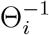, provides a point estimate of the individual-specific covariance matrix, and its inverse Θ_*i*_ serves as the corresponding precision matrix estimate.

#### Lemma 2.1

*Under assumptions* (2) *and* (3), *the posterior variance of the* (*k, l*)*-th entry* 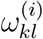

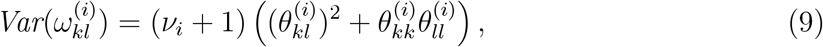

*where* 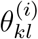*denotes the* (*k, l*)*-th entry of* Θ_*i*_.

*Proof*. Follows from Theorem 2.1 and the moments of Wishart (Mardia et al. 2024).

#### Remark

The partial correlation between variables *k* and *l* for individual *i* is obtained by standardizing the posterior precision estimate:

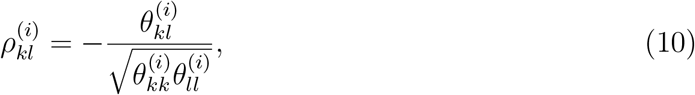

where 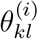denotes the (*k, l*)-th entry of Θ_*i*_.

For high-dimensional omics data where *g* is of the order of thousands, directly computing Θ_*i*_ for each individual requires inverting *n* separate *g* × *g* matrices, which is computationally prohibitive. The following lemma shows that Θ_*i*_ can be expressed entirely in terms of the single population-level matrix inversion *V* ^−1^ and matrix multiplications, reducing *n* matrix inversions to one. This makes SIREN highly scalable to high-dimensional omics data.

#### Lemma 2.2

*By the Sherman-Morrison formula we can show that*

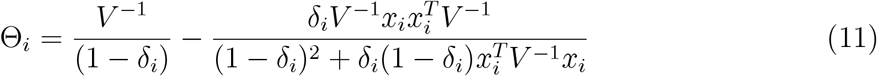

*where* 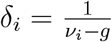 *and V* = (1 − *λ*)*S* + *λT*.

Proof is provided in supplementary material S1.

The parameter *δ*_*i*_ quantifies the relative contribution of two sources of information: the population-level precision *V* ^−1^, estimated from all samples, and the individual-specific centered observation *x*_*i*_. Since *ν*_*i*_ − *g* ≥ 1, we have 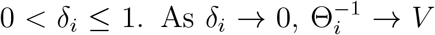, so the individual network collapses to the population average; as 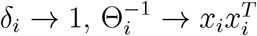, giving full weight to the individual observation. Thus smaller *δ*_*i*_ is appropriate for homogeneous populations where individuals share a common precision structure, while larger *δ*_*i*_ reduces shrinkage bias in heterogeneous populations. In the following section we describe a data-driven approach for calibrating *δ*_*i*_.

### 2.2 Specifying Prior Degrees of Freedom

Since 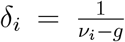 is a one-to-one function of *ν*_*i*_, obtaining a data-driven estimate of *ν*_*i*_ automatically yields an estimate of *δ*_*i*_. By standard properties of the Inverse Wishart distribution (Mardia et al. 2024), under assumption (3), the prior variance of the *k*-th diagonal entry of Σ_*i*_, denoted 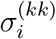, is

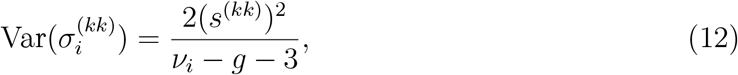

where *s*^(*kk*)^ denotes the *k*-th diagonal entry of *S*. Summing over *k* = 1, …, *g*, we get

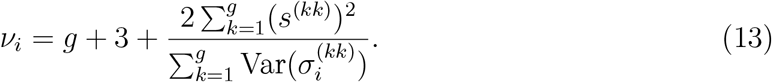

The only unknown on the right-hand side is Var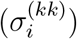, which we estimate via a method-of-moments approach using jackknife resampling. Specifically, we compute *n* leave-one-out estimates of the variance of the *k*-th omics variable, 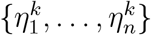, and estimate Var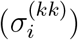 by their sample variance:

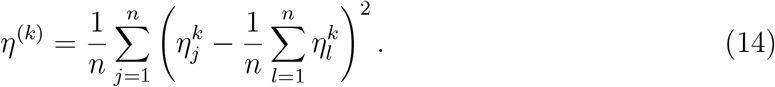

Substituting *η*^(*k*)^ for Var 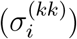 in (13) yields

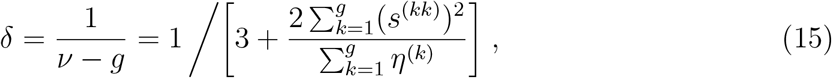

The procedure above yields a common estimate *δ* for all individuals, which is computationally efficient and eliminates subjective hyperparameter specification. When individual-specific *δ*_*i*_ values are preferred, for example in the presence of outliers or samples from rare disease subtypes, Saha et al. (2024b) describe a data-driven leave-one-out procedure that can be adopted for SIREN at the cost of increased computation.

### 2.3 Hypothesis Testing

For notational simplicity, we ignore the individual index *i* from the following discussion. We replace *ν*_*i*_ by *ν*, Ω_*i*_ by Ω and Θ_*i*_ by Θ.

#### Theorem 2.2

*Under the null hypothesis H*_0_ : *θ*_*kl*_ = 0, *the marginal posterior distribution of ω*_*kl*_ *follows a Variance Gamma distribution with mean µ* = 0, *asymmetry β* = 0, *scale parameter* (*θ*_*kk*_*θ*_*ll*_)^1*/*2^*/*(*ν* − *g*) *and shape parameter* (*ν* + 1).

Proof is provided in supplementary material S1.

#### Remark

For large *ν*, the posterior distribution of *ω*_*kl*_ under *H*_0_ is approximately *N* (0, (*ν* +1)*θ*_*kk*_*θ*_*ll*_*/*(*ν* −*g*)^2^) by the normal approximation to the Variance Gamma distribution (Fischer et al. 2025, Section 2.5 and 2.7).

#### Theorem 2.3

*Let* 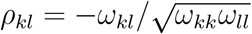*represent the posterior partial correlation between the k-th and l-th molecular variables. The tail probability of the marginal posterior satisfies*

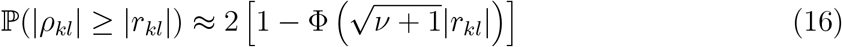

*for large prior degrees of freedom ν, where* Φ(·) *denotes the standard normal CDF*.

Proof is provided in supplementary material S1.

Thus, from the above theorem, we conclude that when the degrees of freedom parameter *ν* is large, the null hypothesis on the partial correlation between the *k*-th and the *l*-th omics variable *H*_0_ : *ρ*_*kl*_ = 0 will have a p-value approximately equal to the right side of (16).

#### Remark

The degrees of freedom *ν > g* + 3 where *g* is the dimensionality of the omics observations, which is typically of the order of thousands in most real data applications (e.g., *g* = 4,003 in the yeast knockout experiment and *g* = 1,981 in the lung adenocarcinoma application), ensuring *ν* is at minimum of the order of thousands and making this approximation theoretically justified in practice.

### 2.4 Connection with James-Stein Type Estimators

James-Stein estimators leverage Stein’s paradox (Stein 1956) by using information from all parameters simultaneously to improve estimates for individual parameters. Suppose we observe *n* ≥ 3 independent samples from *n* populations: *X*_*i*_ ~ *N*(*µ*_*i*_, *σ*^2^), *i* = 1, …, *n*. The usual maximum likelihood estimators 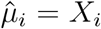are inadmissible under squared error loss when *n* ≥ 3 (James et al. 1961). The James-Stein estimator improves upon these by shrinking towards the overall mean 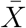 :

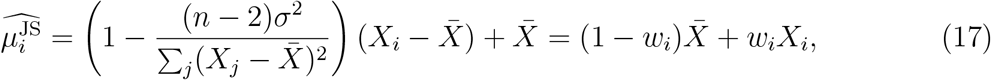

where 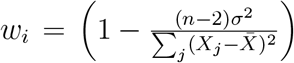. SIREN extends this shrinkage philosophy to individual-specific covariance matrix estimation. The maximum likelihood estimate of Σ_*i*_ based on a single observation is 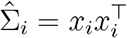, which is rank-deficient when *g >* 1 and inadmissible under squared Frobenius loss. SIREN instead proposes the shrinkage estimator:

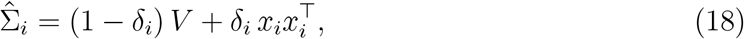

where *V* is the population-level OAS covariance estimate (Chen et al. 2010) and *δ*_*i*_ ∈ (0, 1) is estimated via the empirical Bayes procedure of Section 2.2. This estimator shrinks the rank-1 MLE towards the population average *V*, analogously to how the James-Stein estimator shrinks individual means towards the grand mean. In (18), *V* encodes the population-level covariance structure, while the rank-1 term 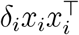 encodes the individual-specific deviation from the population. The estimator (18) coincides with 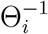, the posterior mean covariance derived in Theorem 2.1, establishing the similarity between the empirical Bayes posterior mean and the James-Stein type shrinkage estimator.

The optimal shrinkage parameter *δ*^∗^ minimizing the average Frobenius risk depends on the unknown Σ_*i*_ and is not directly estimable. Here we adopt an empirical Bayes calibration strategy which estimates *δ* via diagonal variance matching and is computationally efficient. Deriving the SURE-optimal *δ* for individual-specific precision matrix estimation (Stein 1981) represents an interesting direction for future work.

The connection between empirical Bayes and James-Stein shrinkage has been studied for the normal means problem (Efron & Morris 1972*a*). SIREN extends this to individual-specific covariance matrix estimation, building on a rich literature on James-Stein type estimators for population-specific estimates of covariance matrices and their eigenvectors (Haff 1980, Ledoit & Wolf 2004, Chen et al. 2010, Goldberg & Kercheval 2023).

## 3 Applications to Simulated Data

We compare SIREN with three methods for estimating population-specific partial correlation matrices: the Ledoit-Wolf (LW) estimator (Ledoit & Wolf 2004), the graphical LASSO (GL) (Friedman et al. 2008) and the Oracle Approximating Shrinkage (OAS) estimator (Chen et al. 2010). We compare the four methods on four simulation settings: (setups A and B) where the samples come from homogeneous populations where all samples have the same true partial correlation structure and (setups C and D) heterogeneous populations where a certain percentage of samples come from one homogeneous population and the rest of the samples come from another homogeneous population with different true partial correlation structures. Since for multivariate Gaussian random variables the partial correlation matrices can be directly computed from the precision matrix, we specify the structure of the precision matrix in our four simulation setups (A-D). In setups A and C, we assume there is only one omics data type with *p* nodes (molecular variables) and edge density in the partial correlation networks denoted by *η*. In setups B and D, we assume there are two omics layers, with *p*_1_ and *p*_2_ nodes (molecular variables) respectively in the two omics modalities. Let *η*_11_, *η*_22_ and *η*_12_ denote the edge density among nodes within the first layer, among nodes within the second layer and between nodes from the first and the second layer respectively. The four setups are described below:

A. Samples are generated from a homogeneous population where every individual has the exact same precision matrix Ω_1_ with edge density *η* = 0.005.
B. Samples are generated from a homogeneous population where every individual has the exact same precision matrix Ω_2_. Data consists of two omics layers with unequal edge density within and between layers: *η*_11_ = 0.005, *η*_22_ = 0.01 and *η*_12_ = 0.01.
C. Samples are generated from a mixture of two homogeneous populations, where 60% of samples are simulated from a homogeneous population where every individual has the exact same precision matrix Ω_1_, same as in setup A. The other 40% of samples are simulated from a different homogeneous population, whose precision matrix is generated by removing 10% randomly chosen edges from Ω_1_ and adding another 10% randomly chosen non-zero edges in places where no edge existed in Ω_2_.
D. The samples are generated from a mixture of two homogeneous populations, where 60% of samples are simulated from a homogeneous population where every individual has the exact same precision matrix Ω_2_, same as in setup B. The other 40% of samples are simulated from a different homogeneous population, whose precision matrix is generated by removing 10% randomly chosen edges from Ω_2_ and adding another 10% randomly chosen non-zero edges in places where no edge existed in Ω_2_.

The precision matrices Ω_1_ and Ω_2_ are square matrices. To generate them we start with an identity matrix of dimension *p* and randomly replace a proportion *η*_*k,l*_ of zeros on the off-diagonals by uniform random numbers drawn from a uniform distribution ranging between −1 and 1. Then we replace the diagonal entries *ω*_*ii*_ by the sum of the absolute values of all off-diagonal entries in the same row _*j*_ |*ω*_*ij*_|, and add a small positive number *ϵ* = 0.0001 to it. Finally, to normalize the off-diagonal entries *ω*_*ij*_ of the precision matrix, we divide them by 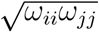so that the resulting matrix resembles a true precision matrix. Next, for each setup (A-D), using the corresponding precision matrices as described above, we simulate individual omics profiles from multivariate Gaussian distributions with mean zero and covariance equal to the inverse of the corresponding precision matrix. To resemble random sampling error that is present in real data, we added small random Gaussian noises from *N* (0, 0.1) to each entry of the simulated data matrix.

For each setup (A-D), we simulate multiple datasets for varying dimensions as well as varying number of samples. For each dataset, we infer sample-specific partial-correlation networks using SIREN and the three population-specific partial correlation networks which assumes that all samples have the exact same network and estimates a single network for all samples. For each sample, we compute the area under the receiver operating characteristic curve (AUC) for each method by comparing the true network edges with the estimated network edges. AUC quantifies the efficacy of each method in distinguishing between true and false interactions, i.e. nonzero and zero network edges respectively. Then for each simulated dataset, we compare the median AUC over all samples.

First, we fix dimensions to *p* = 600 for setups A and C and *p*_1_ = 500, *p*_2_ = 100 for setups B and D, and vary sample size *n* = 100, 500, 1000, 2000 (Figure 1). For homogeneous populations, SIREN achieves comparable performance to LW, GL and OAS despite assuming heterogeneous structure. For heterogeneous populations, SIREN consistently achieves substantially higher AUC, as population-specific methods average out sample-specific heterogeneity while SIREN recovers individual-specific partial correlation structures.

**Figure 1.**
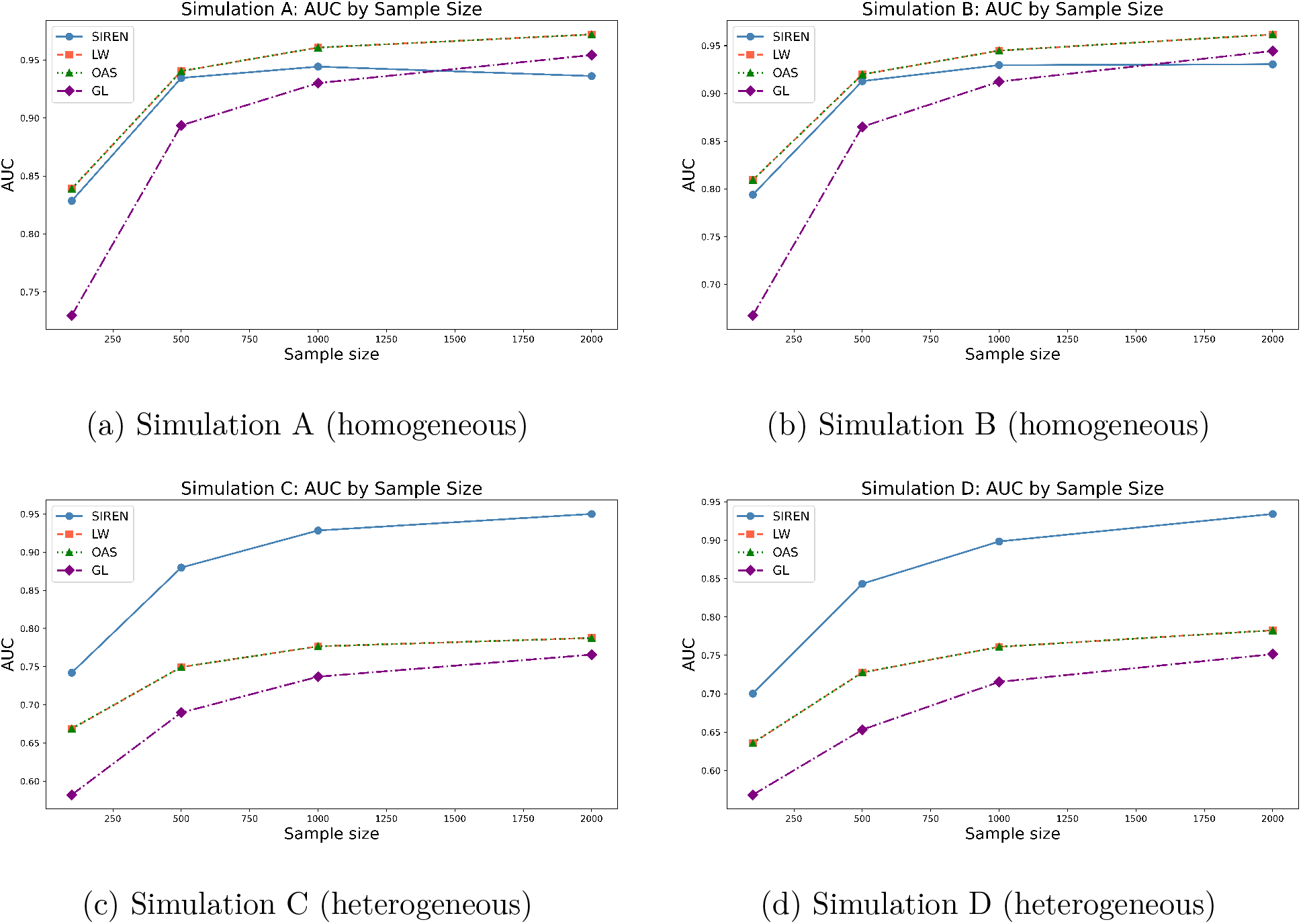
AUC of SIREN vs competing methods as sample size *n* varies. As *n* increases, AUC improves across all settings; SIREN maintains advantage in the heterogeneous settings (C, D), while performance is comparable to competitors in the homogeneous settings (A, B).

Next, we fix *n* = 500 and vary total dimensions *p* = 50, 100, 500, 750, 1000, assigning *p*_1_ = 0.8*p* and *p*_2_ = 0.2*p* for setups B and D (Figure 2). As dimension increases, AUC decreases for all methods. SIREN maintains comparable performance to population-specific methods in homogeneous settings and consistently higher AUC in heterogeneous settings.

**Figure 2.**
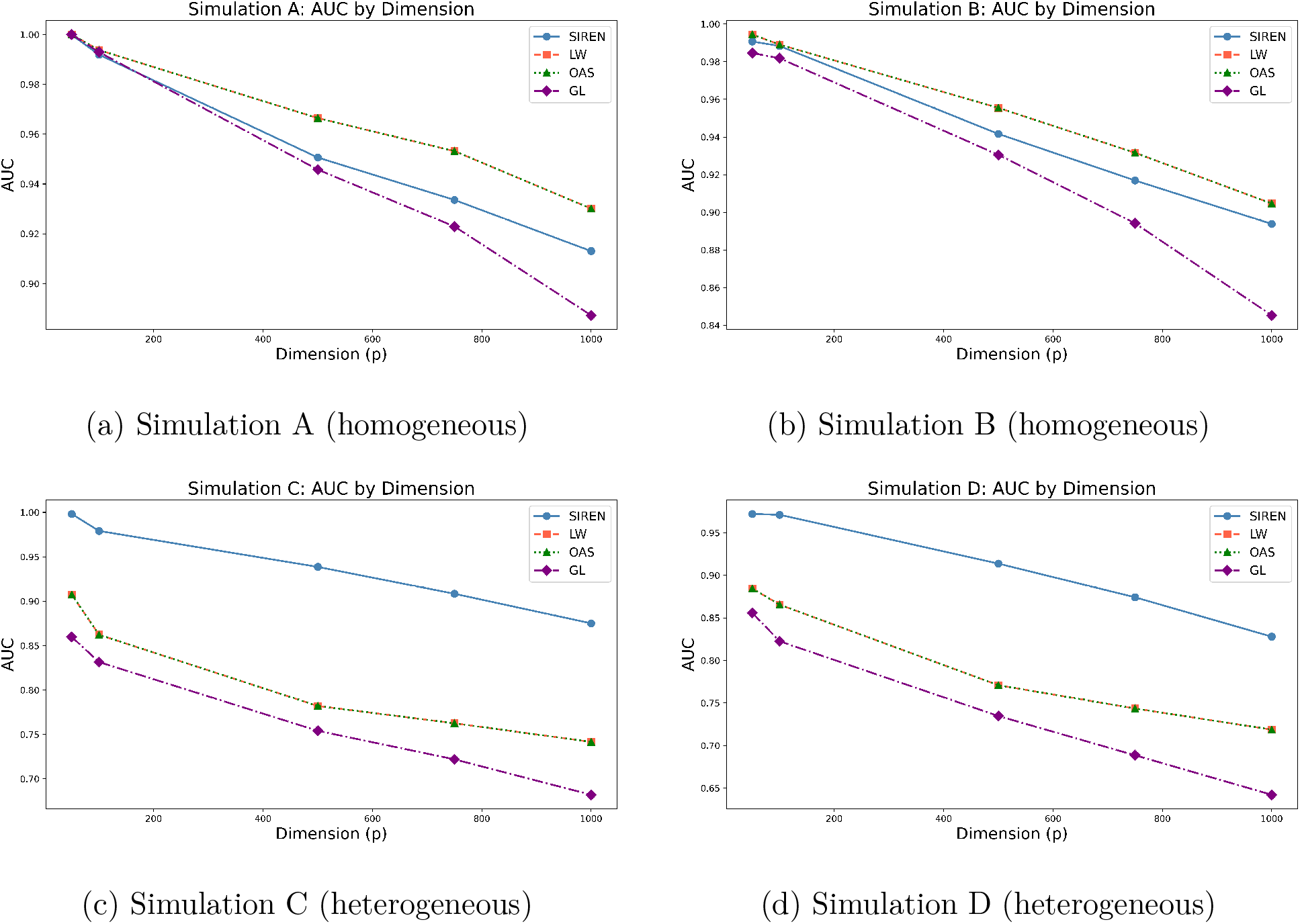
AUC of SIREN vs competing methods as dimension *p* varies. As *p* increases, AUC decreases for all methods; SIREN maintains advantage in the heterogeneous settings (C, D), while performance is comparable to competitors in the homogeneous settings (A, B).

For heterogeneous setups C and D, we additionally vary the minority subpopulation proportion *ρ* = 0.1, 0.2, 0.3, 0.4 (Figure 3). As *ρ* increases, AUC decreases slightly for SIREN but rapidly for other methods. The performance of SIREN compared to other methods gets even better as *ρ* increases, i.e. as the population becomes more heterogeneous.

**Figure 3.**
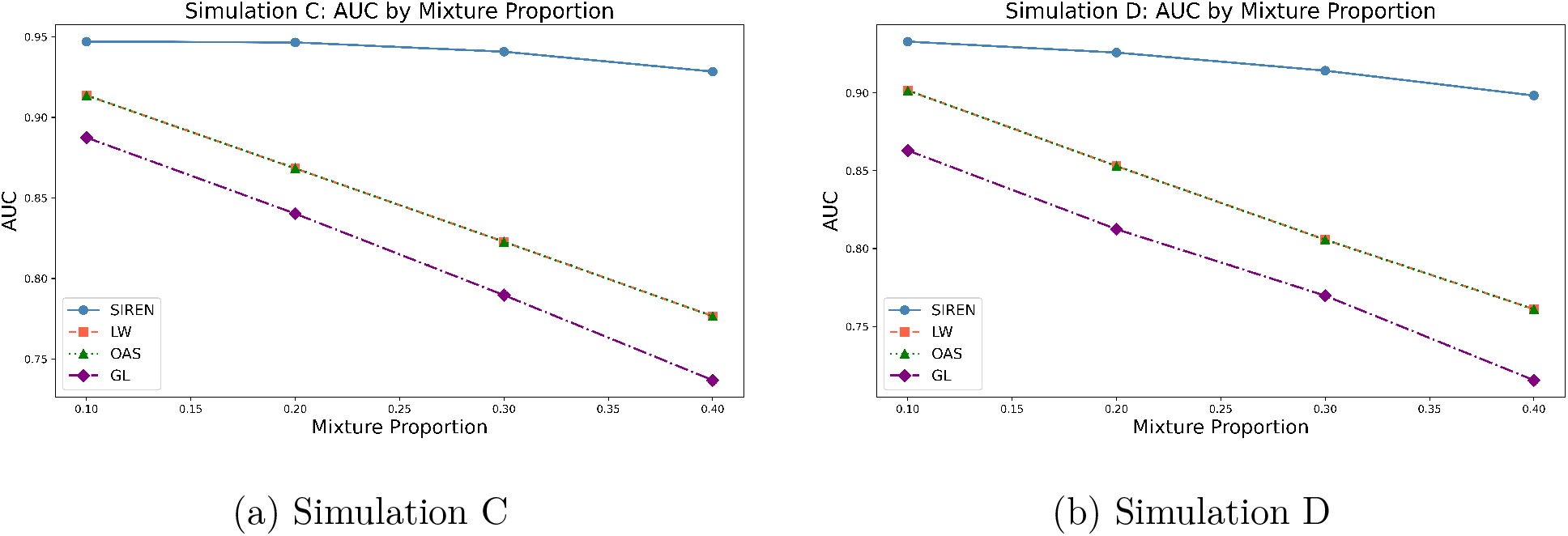
AUC of SIREN vs competing methods as the mixture proportion of the minority subpopulation increases, under heterogeneous simulation settings C and D. As the population becomes more heterogeneous (larger minority proportion), SIREN’s advantage over competitors grows, while competing methods degrade substantially.

Memory usage and compute time are reported in Supplementary Figure S1. For *p* = 1000, a single network estimation takes approximately 0.2 seconds, demonstrating scalability.

## 4 Application to Yeast TF Knockout Data

To validate SIREN’s ability to recover known biology, we applied it to a yeast transcription factor (TF) knockout dataset consisting of 132 pseudobulk samples across 11 TF deletions and 11 growth media conditions (Jackson et al. 2020). Twelve yeast strains including a wild-type control and 11 TF deletions spanning four well-characterized nitrogen-sensing pathways were grown in 11 distinct media conditions with different carbon and nitrogen sources. The knocked-out TFs cover the Nitrogen Catabolite Repression (NCR) pathway (GAT1, GLN3, DAL80, DAL81, DAL82, GZF3), the General Amino Acid Control pathway (GCN4), the Ssy1-Ptr3-Ssy5 amino acid sensing pathway (STP1, STP2), and the Retrograde signaling pathway (RTG1, RTG3), spanning both transcriptional activators and repressors, with several TFs forming known heterodimeric pairs (e.g. RTG1 and RTG3) that serve as internal positive controls. We computed pseudobulk expression profiles by averaging single-cell RNA-sequencing counts across all cells in each genotype-medium combination.

When a TF is knocked out, the biological processes it regulates show altered expression relative to wild-type. By comparing individual-specific network edges between the knocked-out TF and all other genes across knockout and wild-type samples, we can identify pathways showing differential connectivity upon TF deletion. If these pathways correspond to canonical targets of each TF, this provides direct evidence that SIREN recovers true biological signal. Crucially, SIREN was applied without any knowledge of the knockout labels, i.e., the partial correlation networks were estimated from gene expression data alone. We applied SIREN to the top 4,003 most variable genes including all 11 knocked-out TFs, yielding 132 individual-specific partial correlation networks. For each TF, we identified edges differentially connected to the knocked-out TF relative to wild-type, using Wilcoxon rank-sum tests. Then we ranked the genes by the signed Wilcoxon statistic, and performed ranked gene set enrichment analysis using Gene Ontology Biological Process gene sets (Aleksander et al. 2023, Subramanian et al. 2005). We observe that SIREN consistently recovers biologically coherent pathway targets for TF knockouts (Figure 4 and Supplementary Figures S2-S11). GCN4, a master transcriptional activator of amino acid biosynthesis under nutrient starvation (Hinnebusch 2005), showed strong enrichment of cytoplasmic translation, ribosome assembly, and rRNA processing pathways (FDR *<* 0.1), with arginine metabolic process, a canonical GCN4 target (Hinnebusch 2005) also enriched (FDR *<* 0.1), directly validating recovery of known GCN4 biology at the network level. Among the Nitrogen Catabolite Repression (NCR) pathway TFs, GLN3 (activator) and DAL80 (repressor) (Cunningham & Cooper 1991) showed opposite enrichment directions for translation-related pathways. Specifically, GLN3 knockout upregulated translation and ATP biosynthesis while DAL80 knockout suppressed cytoplasmic translation (FDR *<* 0.1), correctly reflecting their antagonistic regulatory roles. DAL81 and DAL82, known to form a functional transcriptional complex (Scott et al. 2000), showed similar pathway enrichment patterns with both exhibiting upregulation of translation, ATP synthesis, and protein folding (FDR *<* 0.1), providing an internal positive control showing that SIREN’s network estimates are biologically coherent. Among the Ssy1-Ptr3-Ssy5 (SPS) pathway TFs, STP1 and STP2 showed divergent enrichment patterns consistent with their known overlapping but functionally distinct roles (Andréasson & Ljungdahl 2002): STP1 knockout upregulated translation and mitochondrial ATP synthesis, while STP2 knockout showed an unexpected suppression of translation and energy metabolism alongside upregulation of double-strand break repair and DNA recombination (FDR *<* 0.1), consistent with their known functionally distinct roles. Finally, the retrograde pathway TFs RTG1 and RTG3, which function as heterodimeric TFs (Komeili et al. 2000) that activates TCA cycle genes in response to mitochondrial dysfunction, both showed suppression of glyoxylate metabolism, pyruvate metabolism, and TCA cycle-related processes (FDR *<* 0.1), validating their known role in retrograde metabolic signaling.

**Figure 4.**
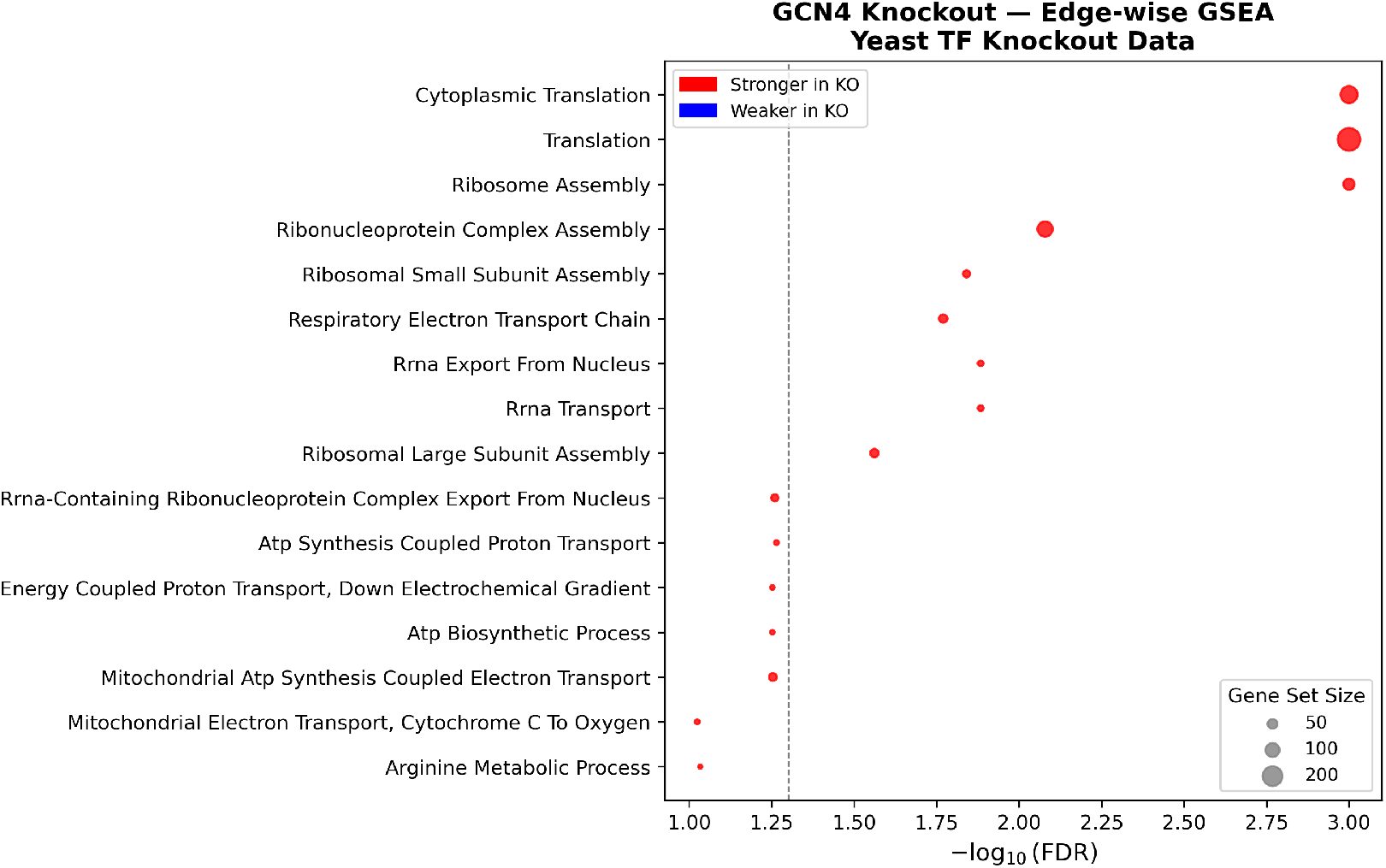
Gene set enrichment analysis of edges differentially connected to GCN4 in GCN4 knockout versus wild-type yeast samples. Bubble color indicates whether edges are stronger (red) or weaker (blue) in the knockout relative to wild-type. Bubble size is proportional to gene set size. Only pathways with FDR *<* 0.1 are shown; the dashed vertical line marks FDR = 0.05. Results for all 11 TFs are shown in Supplementary Figures S2-S11.

Taken together, these results demonstrate that SIREN consistently recovers biologically meaningful pathway-level signatures across four distinct regulatory pathways, correctly reflecting known activator/repressor relationships, heterodimeric partnerships, and pathway-specific metabolic functions. Importantly, these results were obtained without any prior knowledge of genotype labels, demonstrating that individual-specific partial correlation networks estimated by SIREN capture genuine regulatory variation across samples.

## 5 Application to Lung Adenocarcinoma

Lung adenocarcinoma, the most prevalent subtype of lung cancer is the leading cause of cancer-related mortality worldwide (Kanchustambham & Sharma 2026). Despite advances in staging and treatment, patients diagnosed at identical tumor stages often have markedly different survival outcomes, implying that there exists individual-specific differences in molecular regulatory mechanisms of disease that drive heterogeneity in prognosis. Epigenetic dysregulation through DNA methylation is a particularly compelling candidate as promoter methylation can silence tumor suppressor genes or activate oncogenes and these epigenetic states vary substantially across individuals (Gimeno-Valiente et al. 2025).

Identifying which genes are associated with survival through analyzing gene expression or DNA methylation data alone reveals *which* molecular variables matters, but cannot identify whether a gene is prognostically relevant because its expression is epigenetically regulated through methylation, and whether this regulatory relationship varies across patients. Population-level GGM methods assign identical edge weights to every patient and therefore cannot capture inter-patient regulatory heterogeneity or be used directly in survival models. Individual-specific networks address this gap: a survival-associated methylation-expression edge identifies not only that a gene matters for prognosis, but that its expression is driven by epigenetic regulation in a patient-specific manner. Such mechanistic insights have direct therapeutic implications as genes whose survival association operates through methylation-driven regulation are natural candidates for epigenetic therapies such as DNA methyltransferase or histone deacetylase inhibitors (Bates 2020), which cannot be identified from survival analyses on expression or methylation data alone.

We applied SIREN to paired gene expression and DNA methylation data from The Cancer Genome Atlas (TCGA) Lung Adenocarcinoma (LUAD) cohort (Network et al. 2014). We obtained matched RNA-seq and 450K methylation array data for *n* = 448 patients with available survival information. For methylation, we computed M-values (log2 ratio of methylated to unmethylated signal intensities), restricted to CpG probes in promoter regions (TSS200, TSS1500, and 5’UTR), and aggregated to the gene level by averaging M-values across all promoter probes mapping to each gene. To retain biologically relevant features, we performed survival-guided prescreening. We fit a univariate Cox proportional hazards model for each expression gene and each methylation feature separately, adjusting for age, sex, race, smoking history, and tumor stage, and retained features with nominal *p <* 0.01. This yielded *p*_1_ = 1,760 expression genes and *p*_2_ = 221 methylation features (*g* = 1,981 total), ensuring that SIREN focuses on features with marginal evidence of survival association rather than expression variability alone.

We apply SIREN to estimate individual-specific gene-methylation partial correlation networks for each of the *n* = 448 LUAD patients, yielding 1,961,190 individual-specific edge weights per patient. Since population-level methods assign identical edge weights to every patient, they cannot be used as individual-level covariates in survival analyses. SIREN’s individual-specific networks, by contrast, provide patient-level edge weights that capture heterogeneity in gene-methylation regulatory relationships. We focus on the 388,960 expression-methylation edges, as these capture epigenetic regulation of gene expression, which is the primary biological mechanism of interest. For each (expression gene, methylation feature) pair, we fit an edge-wise Cox proportional hazards model, with the individual-specific edge weight as the primary covariate, adjusting for age, sex, race, smoking history and tumor stage. Multiple testing correction is performed using the Benjamini-Hochberg procedure (Benjamini & Hochberg 1995).

We visualize the top survival associated edges identified by the cox model as a network (Figure 5), with edges colored red for hazardous associations (HR *>* 1) and blue for protective associations (HR *<* 1). To further characterize these associations, we stratified patients by median individual-specific edge weight into high and low groups and plotted Kaplan-Meier survival curves separately for early (Stage I-II) and late (Stage III-IV) disease stage (Figure 6). The curves demonstrate that the high and low edge weight groups are consistently separated across both disease stages, suggesting that individual differences in methylation-driven gene regulation stratify patients by survival, independently of tumor stage. Six additional Kaplan-Meier curves are shown in Supplementary Figure S13.

**Figure 5.**
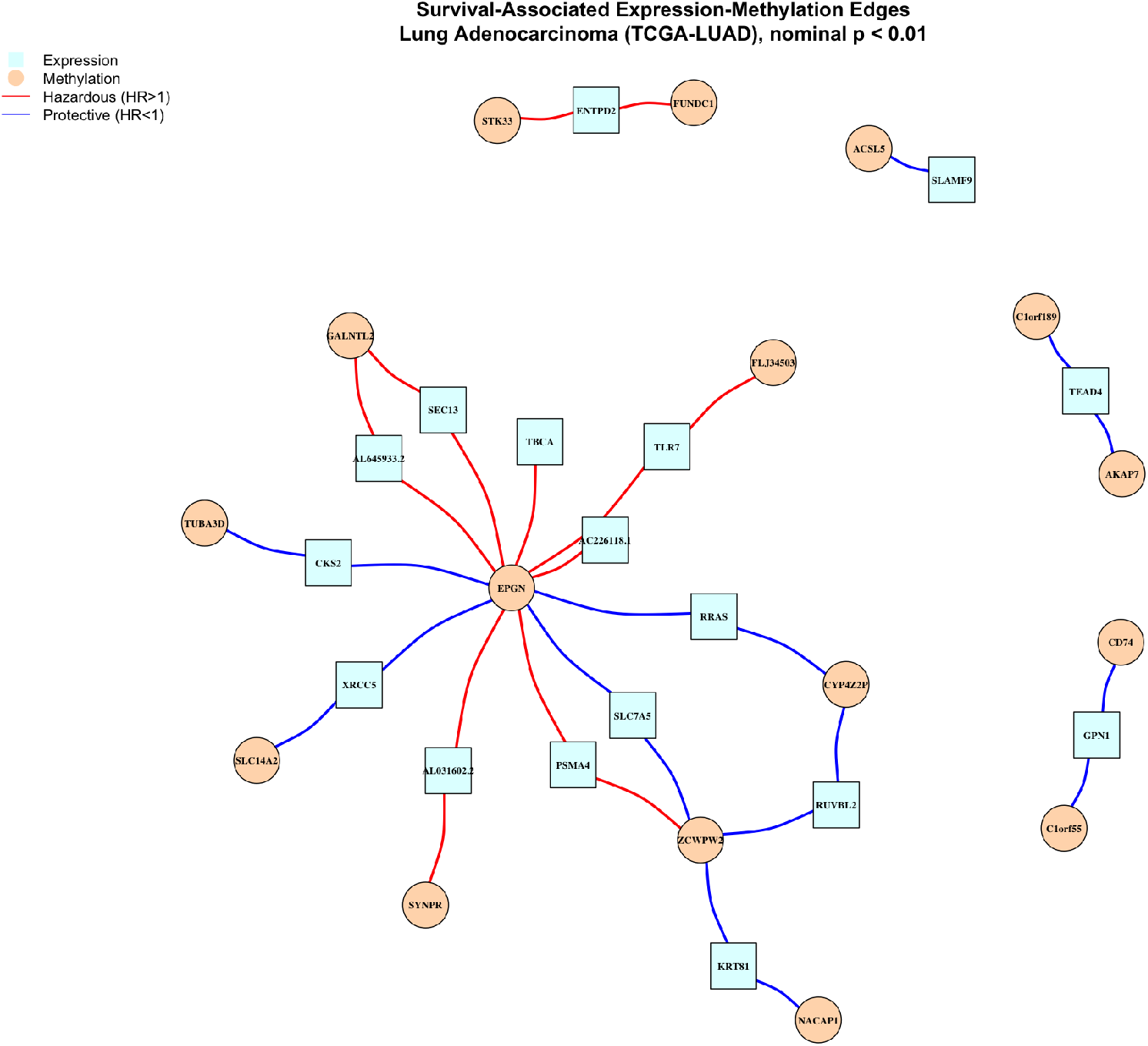
Survival-associated expression-methylation network in lung adenocarcinoma. Each node represents a gene, with expression nodes shown as cyan squares and methylation nodes shown as orange circles. Edges connect expression and methylation features whose individual-specific partial correlation is associated with overall survival (nominal p *<* 0.01). Edge color indicates the direction of association: red edges indicate that higher partial correlation is associated with worse survival (HR *>* 1), while blue edges indicate a protective association (HR *<* 1). Among the top 50 most significant edges, at most two edges per expression gene are shown, yielding 31 edges in total.

**Figure 6.**
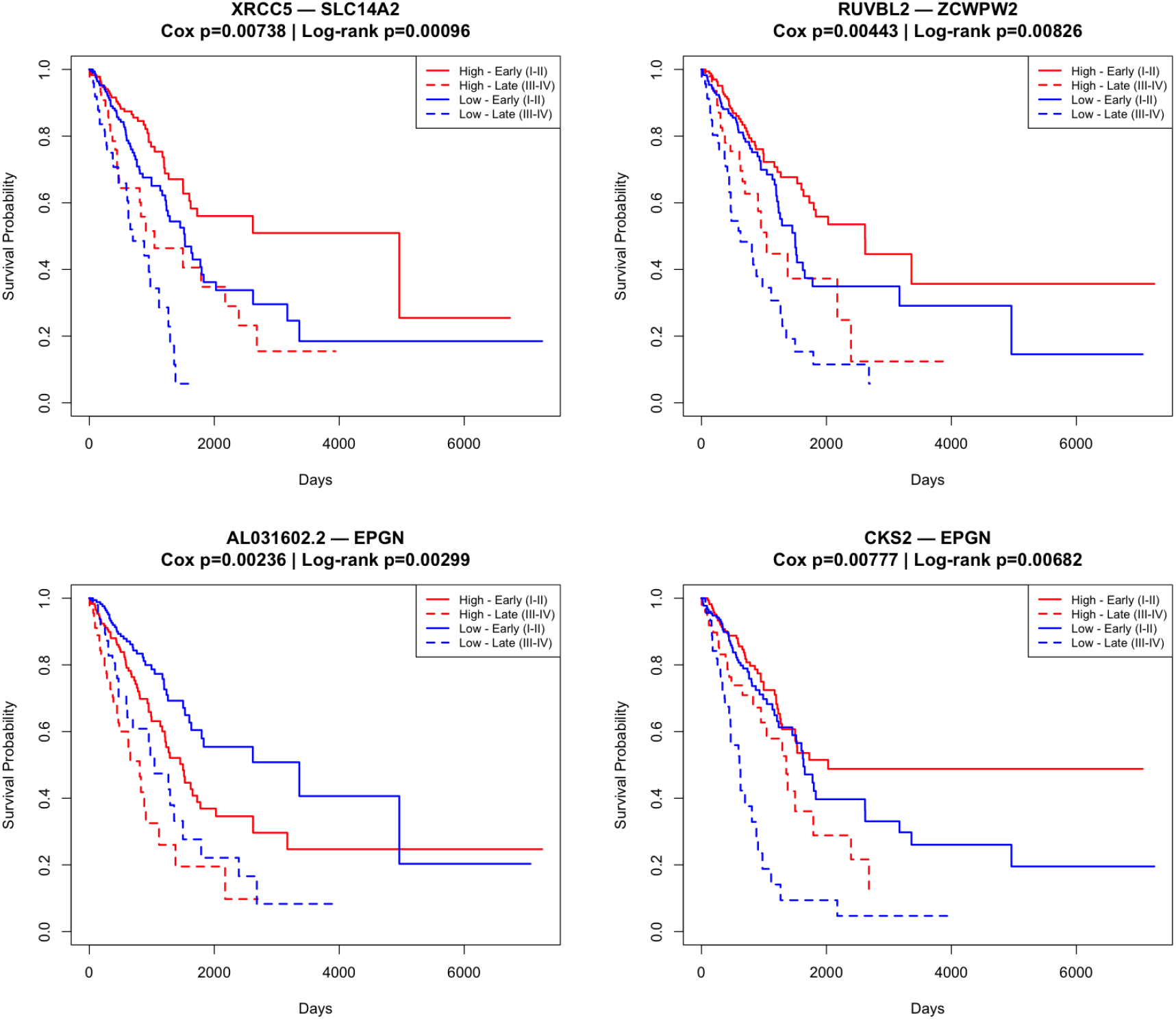
Kaplan-Meier survival curves for four representative survival-associated expression-methylation edges in lung adenocarcinoma. Edge labels denote expression gene followed by methylation gene (e.g., XRCC5–SLC14A2). For each edge, patients are stratified by the median individual-specific partial correlation weight into high (red) and low (blue) groups, with solid and dashed lines representing early (Stage I-II) and late (Stage III-IV) disease respectively. Cox model adjusted for age, sex, tumor stage, smoking status, and race. Log-rank p-values are stratified by tumor stage.

Several biologically relevant genes appear among the top survival-associated edges. *HMGA2*, known to promote LUAD metastasis through PI3K/AKT/VEGFA signaling (Kanchustamb-ham & Sharma 2026), showed a protective survival-associated edge with methylation of *OPHN1. XRCC5*, a key mediator of DNA double-strand break repair (Zhang et al. 2021), showed a protective edge with methylation of *SLC14A2. RUVBL2*, which promotes tumor metastasis through MAPK and Notch signaling (Huang et al. 2025), showed a protective edge with methylation of *ZCWPW2*, consistent with chromatin remodeling as a protective mechanism identified in the pathway enrichment analysis below. *CKS2*, whose overexpression correlates with poor prognosis in LUAD (Wang et al. 2021), showed a protective edge with methylation of *EPGN*, particularly among late-stage patients. Notably, *EPGN* methylation appears as a regulatory hub, with its methylation associated with the expression of multiple survival-relevant genes. Conversely, higher partial correlation between *PBK* expression and methylation of *CLDN16* was associated with worse survival (Ma et al. 2023), consistent with PBK’s known oncogenic role in LUAD.

To identify biological pathways underlying survival-associated epigenetic regulation, we computed for each patient *i* and expression gene *g* the methylation-specific in-degree: 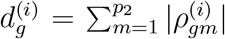, where 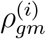 denotes the individual-specific partial correlation between gene *g* and methylation feature *m*. This quantity captures the extent to which gene *g*’s expression is epigenetically regulated through methylation in patient *i*. We fit a gene-wise Cox model with 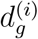 as the covariate, adjusting for age, sex, race, smoking, and tumor stage, ranked genes by signed log p-value, and performed gene set enrichment analysis against Gene Ontology Biological Process gene sets (Aleksander et al. 2023).

We identified 25 significant pathways at *p <* 0.01 (Figure 7). Among these, pathways related to chromatin organization, chromatin remodeling, and epigenetic regulation of gene expression were strongly protective, directly validating the biological relevance of methylation-expression regulatory networks estimated by SIREN and consistent with aberrant promoter methylation as a driver of tumor suppressor silencing in LUAD (Bates 2020). Multiple WNT signaling pathways were also protective, consistent with known tumor suppressive roles of WNT activation in LUAD (Stewart 2014). Stem cell differentiation and regulation of cell differentiation pathways were additionally protective, suggesting that genes controlling cellular differentiation state contribute to LUAD survival heterogeneity. Conversely, axon development was the only hazardous pathway (*p <* 0.01), consistent with emerging roles of neuronal signaling programs in LUAD progression (Mi et al. 2025), though this finding did not survive FDR correction and should be interpreted with caution.

**Figure 7.**
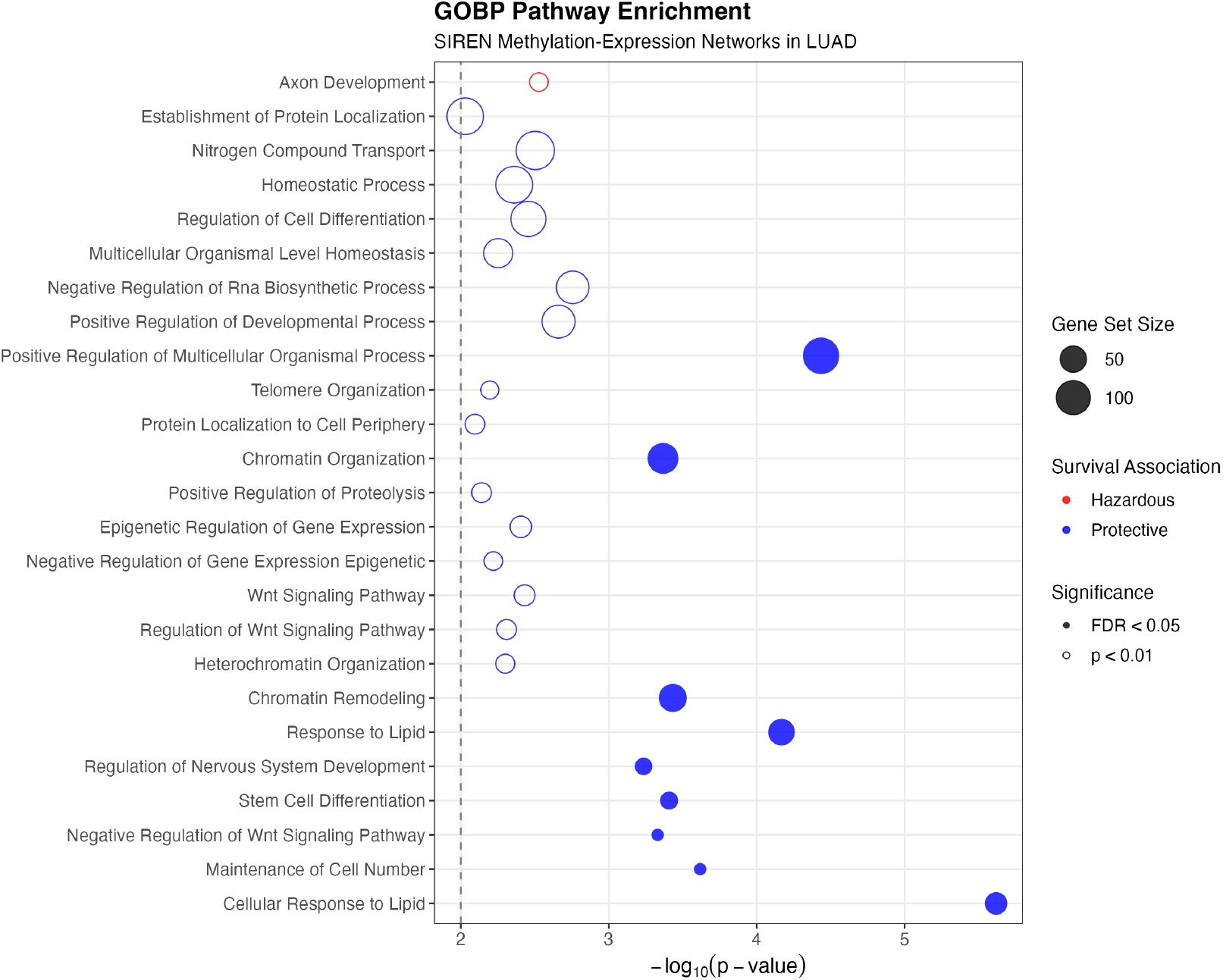
Gene set enrichment analysis of survival-associated expression-methylation edges in lung adenocarcinoma. Bubble plot shows Gene Ontology Biological Process (GOBP) pathways enriched among genes ranked by signed log p-value from the edge-wise Cox model. Only pathways with nominal p-value *<* 0.01 are shown. Bubble color indicates the direction of enrichment: blue bubbles represent pathways negatively associated with hazard (protective), and red bubbles represent pathways positively associated with hazard. Bubble size is proportional to the number of genes in the pathway. Filled circles indicate pathways significant at FDR *<* 0.1. The dashed vertical line marks nominal p-value = 0.01.

## 6 Discussion

We introduce a method for estimating individual sample-specific partial correlation networks through the empirical Bayes principle, deriving Gaussian graphical models for heterogeneous populations. Since sample-specific covariance is not invertible when data dimension exceeds sample size, we use the OAS shrinkage estimator of Chen et al. (2010) as the prior mean of the sample-specific covariance matrices for each sample. Through the use of conjugate priors, we ensure that the posterior distribution of sample-specific precision matrices have closed-form expressions for each sample, thereby alleviating the need for expensive MCMC sampling, thus making our method scalable to very high dimensional data. The resulting posterior mean covariance is a weighted combination of the rank-1 sample-specific update 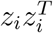, which is the MLE of sample-specific covariance and the population-level OAS estimate *V*. In this spirit our estimator can be thought of as a James-Stein-type estimator for sample covariance for heterogeneous populations. While SIREN is designed for settings where individual-level heterogeneity is of primary interest, it is complementary to mixture GGM methods (Hao et al. 2018, Wu et al. 2024) that estimate group-level networks for predefined or latent subpopulations. In practice, these approaches address different scientific questions and can be used in conjunction: mixture methods to identify subpopulation structure, and SIREN to characterize individual-level variation within and across subpopulations.

The lung adenocarcinoma application demonstrates a key advantage of individual-specific network estimation that population-level methods cannot provide: by assigning each patient a unique set of methylation-expression edge weights, SIREN enables these weights to serve directly as covariates in survival models, revealing how patient-level heterogeneity in epigenetic gene regulation drives differential survival outcomes. The survival-associated edges identified by SIREN implicate chromatin remodeling, WNT signaling, and stem cell differentiation as protective mechanisms in LUAD, suggesting that individual differences in methylation-driven regulation of genes such as *HMGA2, XRCC5*, and *RUVBL2* contribute to the survival heterogeneity observed among patients diagnosed at the same tumor stage. These findings highlight the potential of individual-specific network estimation as a tool for precision oncology, where characterizing patient-level regulatory heterogeneity may identify candidates for epigenetic therapies tailored to individual molecular profiles. More broadly, SIREN is applicable to any setting where individual-level network heterogeneity is of scientific interest, including neuroscience (Gordon et al. 2017) and the molecular hallmarks of aging (Saha et al. 2025, Saha 2026).

A potential limitation of the proposed empirical Bayes framework is its tendency to shrink all sample-specific covariance matrices toward the common prior mean *V*, which is estimated from the full sample. For individuals whose covariance structure deviates substantially from the population average, such as those with rare disease subtypes, this shrinkage may be overly aggressive. This is analogous to the well-known behavior of James-Stein type estimators, which achieve superior average risk across the group at the potential cost of individual unusual cases (Efron & Hastie 2021). A limited translation correction (Efron & Morris 1972*b*), which restricts shrinkage to at most *D* standard deviations away from the individual’s own estimate, could mitigate this effect, and represents a natural direction for future work. Another limitation is that for multi-omics data, a single shrinkage parameter *λ* is applied uniformly across all omics modalities, treating intra-modality and inter-modality covariances equally. In practice, different omics layers may exhibit substantially different levels of within-layer and between-layer correlation structure. A natural extension would adopt modality-specific shrinkage parameters, shrinking each submatrix of the population covariance estimate by a different degree. Finally, the empirical Bayes calibration of *δ* adopted here is a method of-moments estimator rather than direct minimization of the Frobenius risk. Deriving a SURE-optimal estimate of *δ* for individual-specific precision matrix estimation represents an important theoretical direction for future work. This limitation is particularly relevant in cancer applications where tumor heterogeneity may result in a small subgroup of patients with regulatory networks that differ substantially from the population average, and where aggressive shrinkage toward the population mean could obscure clinically important individual differences.

## 8 Data Availability Statement

SIREN is available with the open-source Python package SIGMAnet (https://github.com/Enakshi-Saha/SIGMAnet). The yeast data is publicly available through Gene Expression Omnibus (GEO) with accession number GSE125162. TCGA data is available through the GDC Data Portal. Code and processed data are provided as supplementary files.

## SUPPLEMENTARY MATERIAL

### S1 Proofs of Main Results

#### Proof of Theorem 2.1

From the standard properties of the Wishart distribution, (Mardia et al. 2024), we know that if a symmetric positive definite matrix *M* ~ Wishart(*M*_0_, *ν*), then *M* ^−1^ ~ InverseWishart 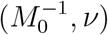. Therefore, the second part of the theorem is equivalent to proving

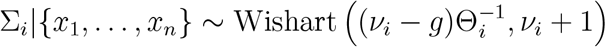

Under the assumption 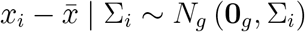, the conditional likelihood of the centered expression of the *i*-th sample can be written as

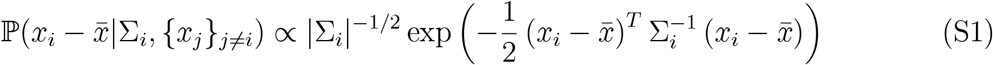

Under assumption Σ_*i*_ ~ IW ((*ν*_*i*_ − *g* − 1)*V, ν*_*i*_), given all other samples in the data, the prior probability density of the covariance matrix of the *i*-th sample can be written as

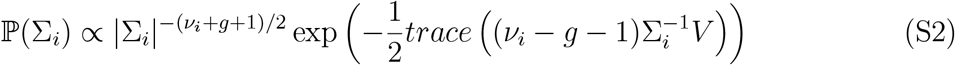

Combining S1 and S2, the posterior distribution of Σ_*i*_ can be written as

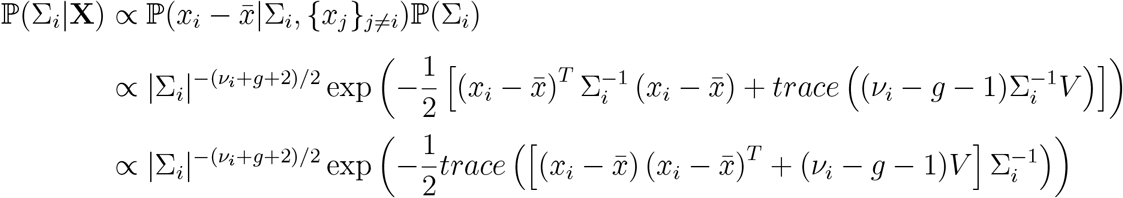

Recognizing that 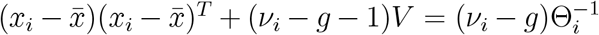, the posterior distribution of Σ_*i*_ is InvWishart 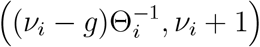, completing the proof.

#### Proof of Lemma 2.1

We use the Sherman-Morrison formula (Sherman & Morrison 1950), which states that given an invertible matrix *A* and two vectors *u* and *v*, the inverse of the sum of *A* and the outer product *uv*^*T*^ is 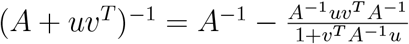. Plugging in *A* = (1 − *δ*_i_)*V* and 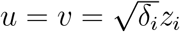 in the formula, we get

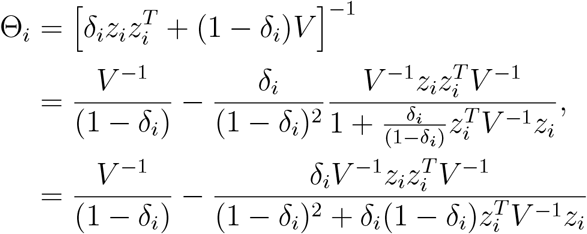

#### Proof of Theorem 2.2

The proof follows from section 4.1.2 of Fischer et al. (2025), which states that the marginal distribution of the off-diagonal entries of a Wishart distribution *W* (Σ, *n*) follows a Variance Gamma distribution 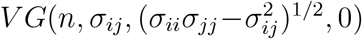. Using (**??**), we get, *ω*_*kl*_ ~ VarGamma 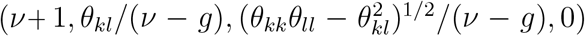. Under *H*_0_ : *θ*_*kl*_ = 0, this simplifies to *ω*_*kl*_ ~ VarGamma(*ν* + 1, 0, (*θ*_*kk*_*θ*_*ll*_)^1*/*2^*/*(*ν* − *g*), 0), giving the result.

#### Proof of Theorem 2.3

From Theorem 2.2, the marginal posterior of *ω*_*kl*_ under *H*_0_ : *θ*_*kl*_ = 0 satisfies

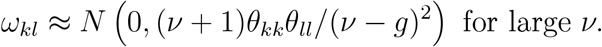

From standard properties of the Wishart distribution (Mardia et al. 2024), 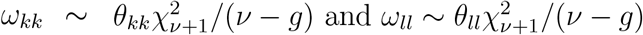. So, for large *ν*,

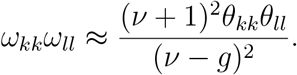

Substituting into 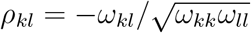 gives

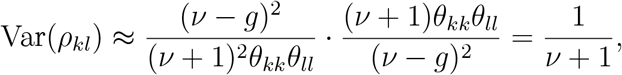

so *ρ*_*kl*_ ≈ *N* (0, 1*/*(*ν* + 1)), and therefore

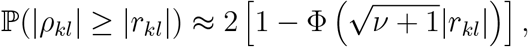

completing the proof.

### S2 Memory Usage and Computational Scalability

Figure S1 reports peak memory usage and per-sample runtime for SIREN across all four simulation setups (A-D). Memory was recorded with varying dimension for fixed sample size *n* = 1000 (Figure S1a) and with varying sample size for fixed *g* = 600 (Figure S1b). Runtime was recorded as the average computation time in seconds for estimating a single sample-specific partial correlation network, fixing *n* = 1000 and varying total dimensions *p* = 50, 100, 500, 750 and 1000. (Figure S1c).

**Figure S1.**
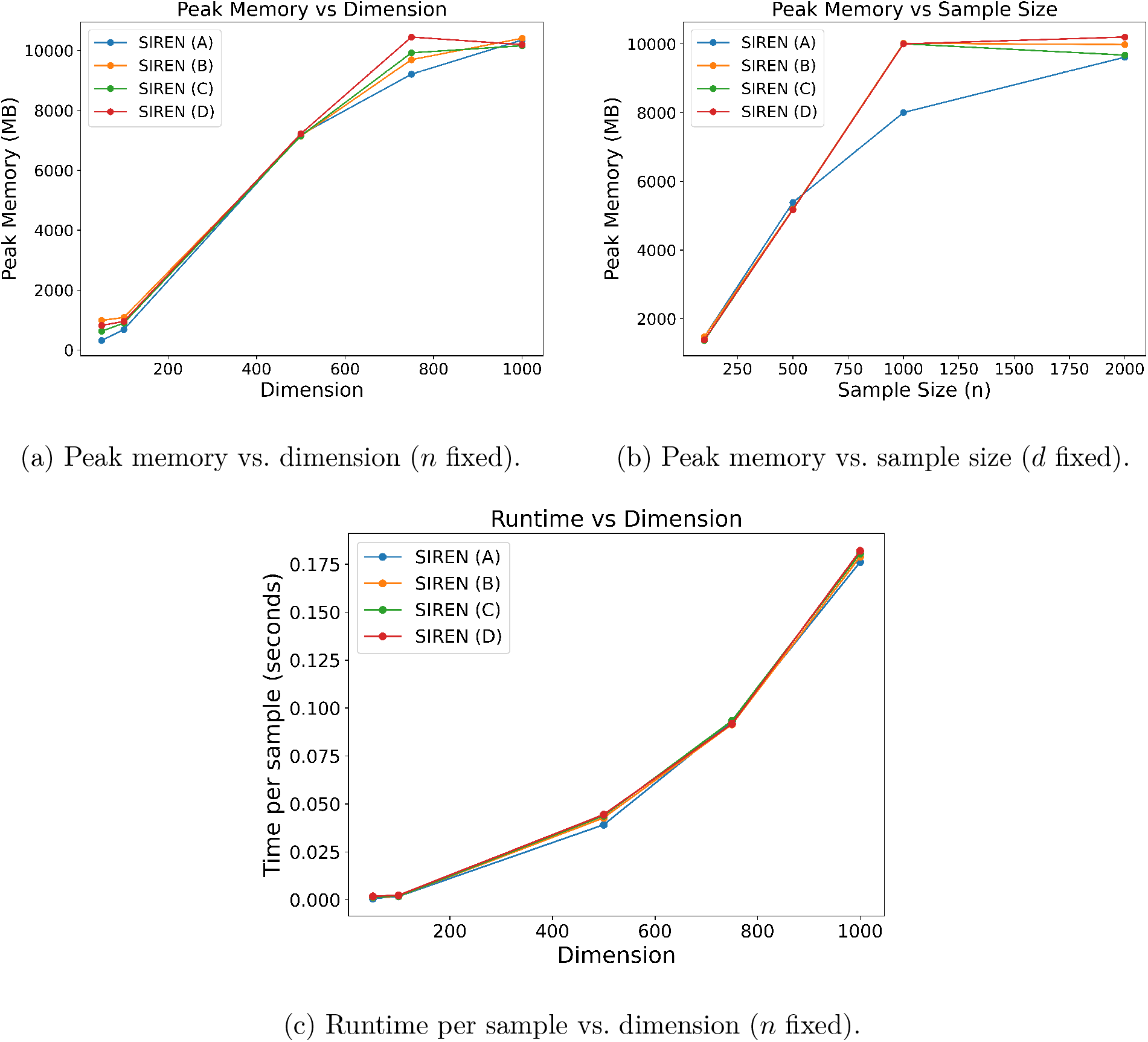
Computational scalability of SIREN across simulations (setups A-D). For all examples, runtime scales quadratically with dimension, consistent with *O*(*g*^2^) complexity of the OAS estimator and Sherman-Morrison rank-1 update. Peak memory plateaus for large *n*, indicating that storage is dominated by *g* × *g* matrix terms rather than sample size.

### S3 Supplementary Figures for Yeast Transcription Factor Knockout Data

**Figure S2.**
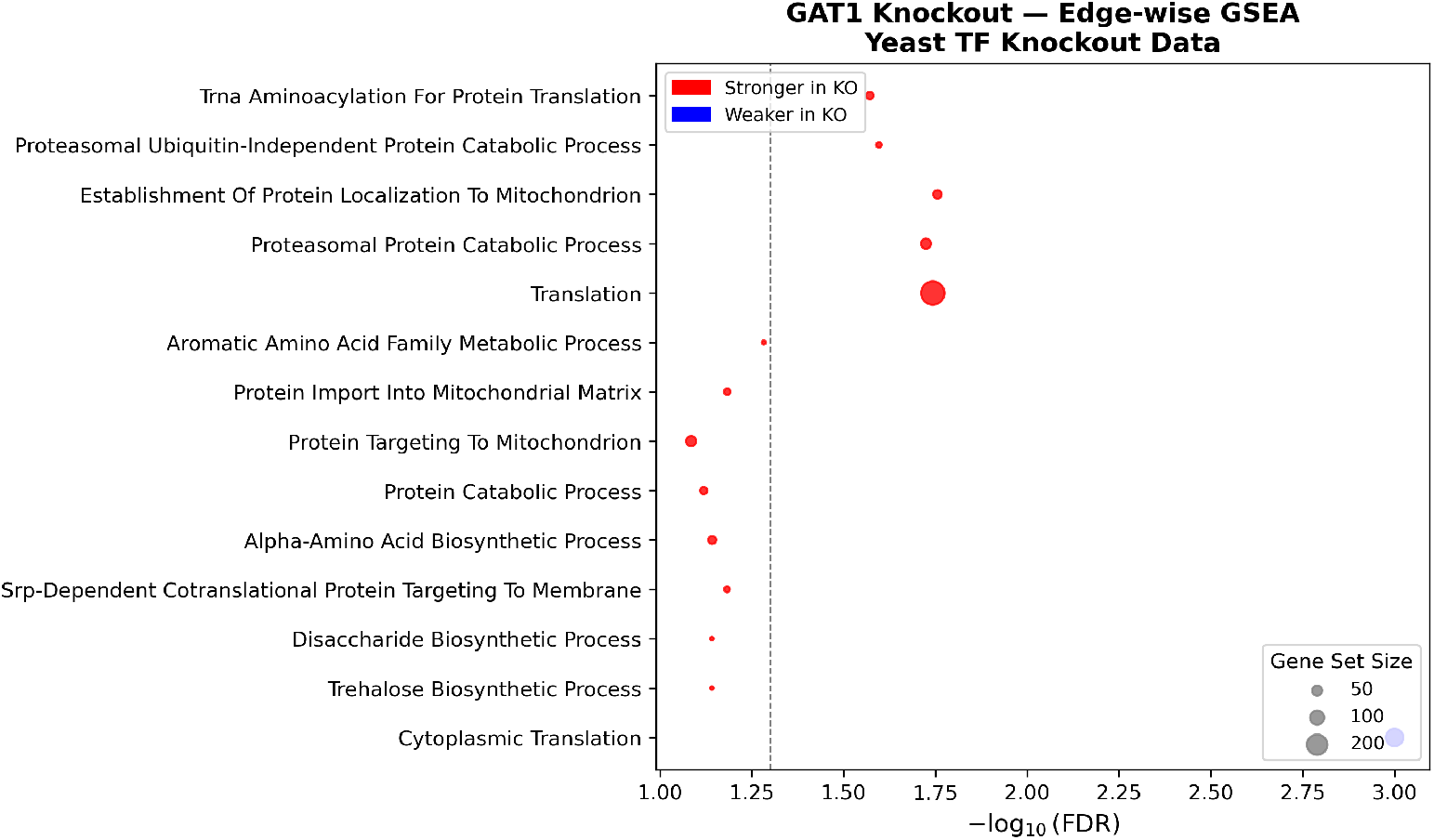
Gene set enrichment analysis of edges differentially connected to GAT1 in GAT1 knockout versus wild-type yeast samples. Bubble color indicates whether edges are stronger (red) or weaker (blue) in the knockout relative to wild-type. Bubble size is proportional to gene set size. The dashed vertical line marks FDR = 0.05. Only pathways with FDR *<* 0.1 are shown.

**Figure S3.**
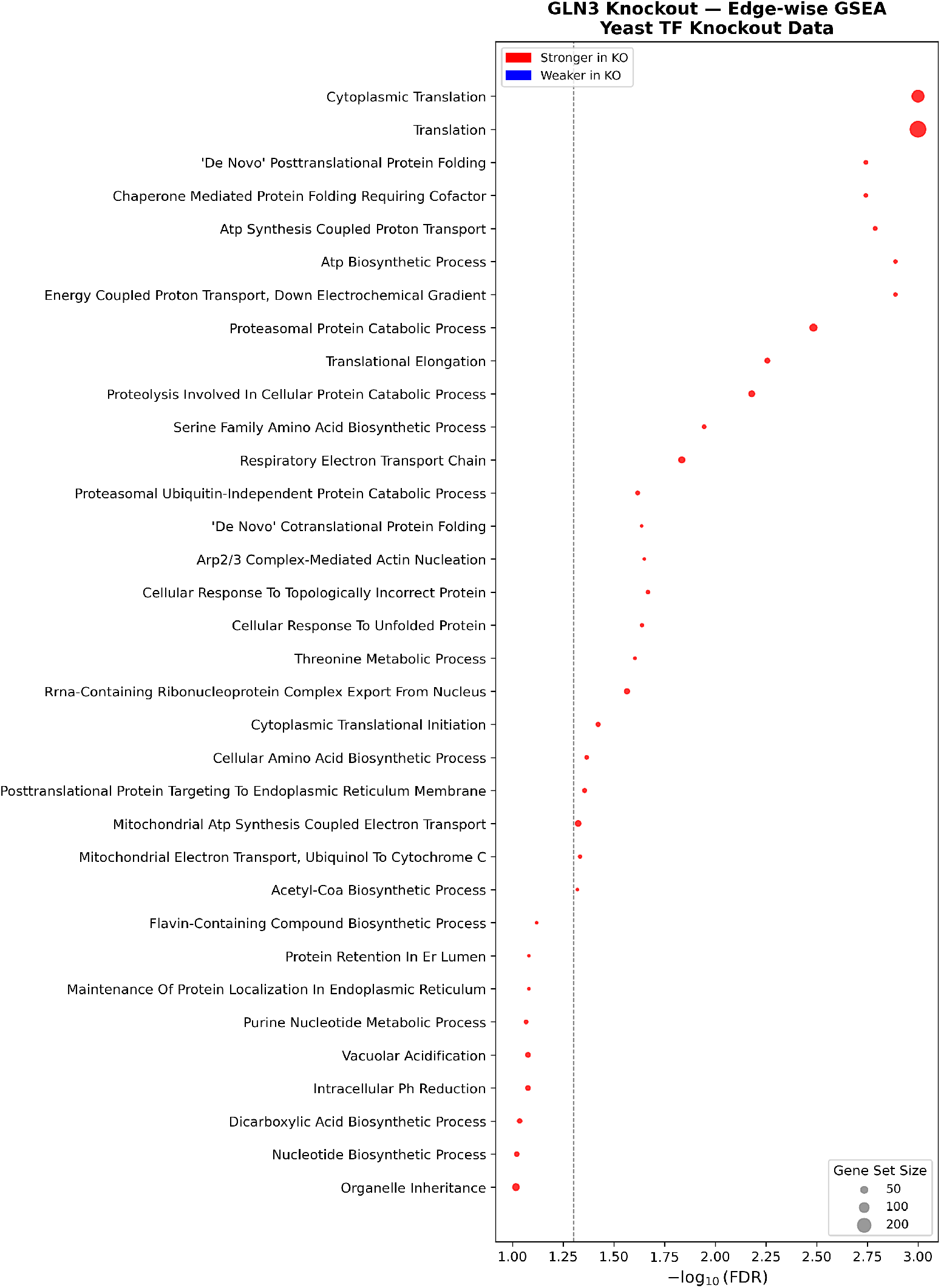
Gene set enrichment analysis of edges differentially connected to GLN3 in GLN3 knockout versus wild-type yeast samples. See Figure S2 for details.

**Figure S4.**
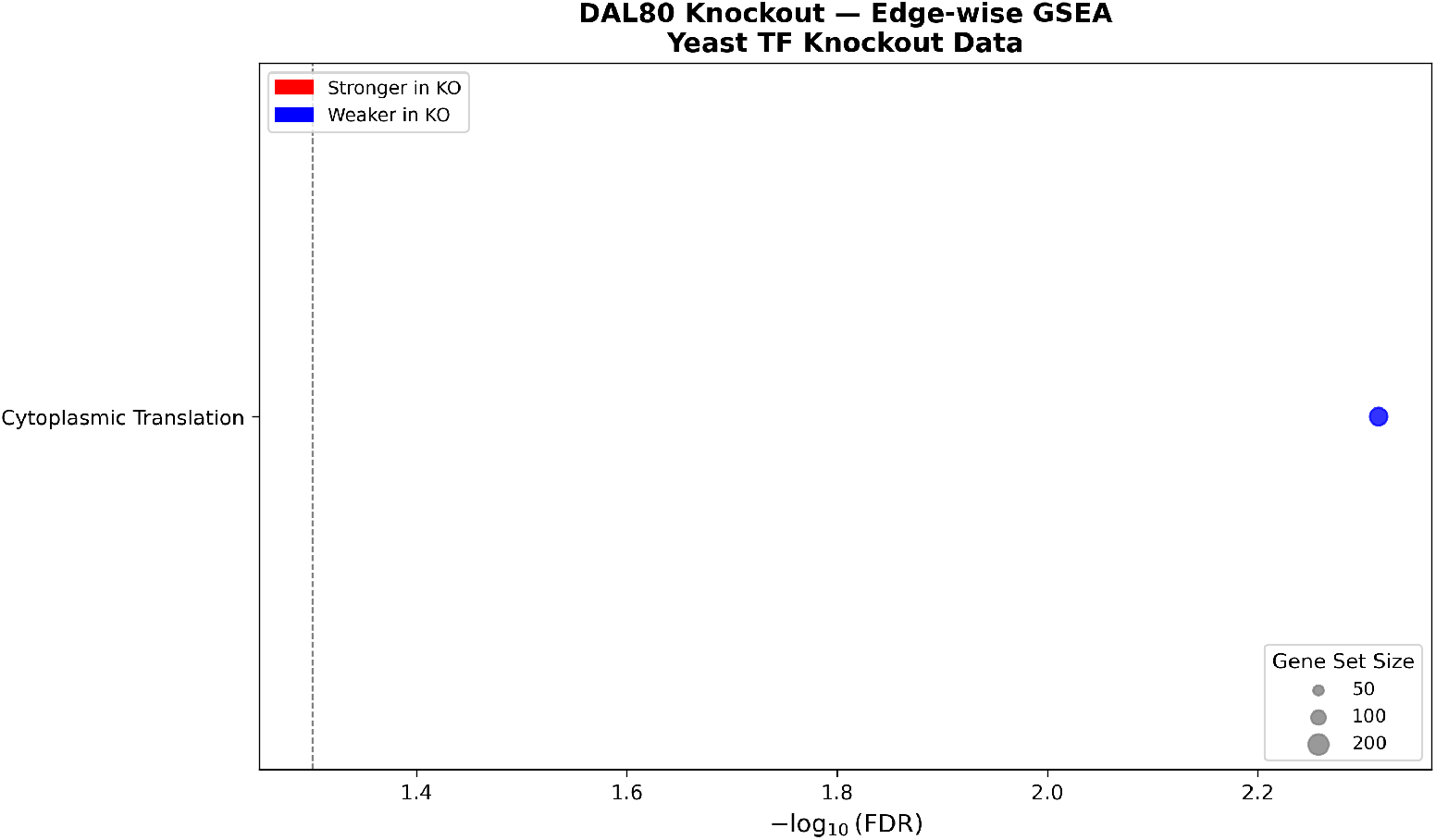
Gene set enrichment analysis of edges differentially connected to DAL80 in DAL80 knockout versus wild-type yeast samples. See Figure S2 for details.

**Figure S5.**
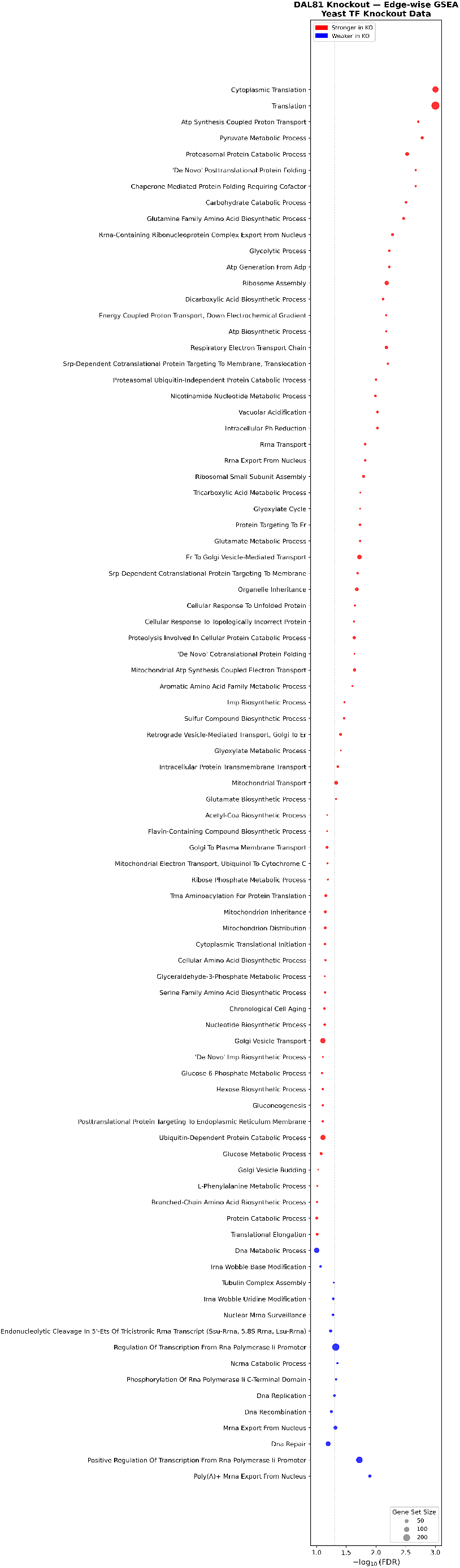
Gene set enrichment analysis of edges differentially connected to DAL81 in DAL81 knockout versus wild-type yeast samples. See Figure S2 for details.

**Figure S6.**
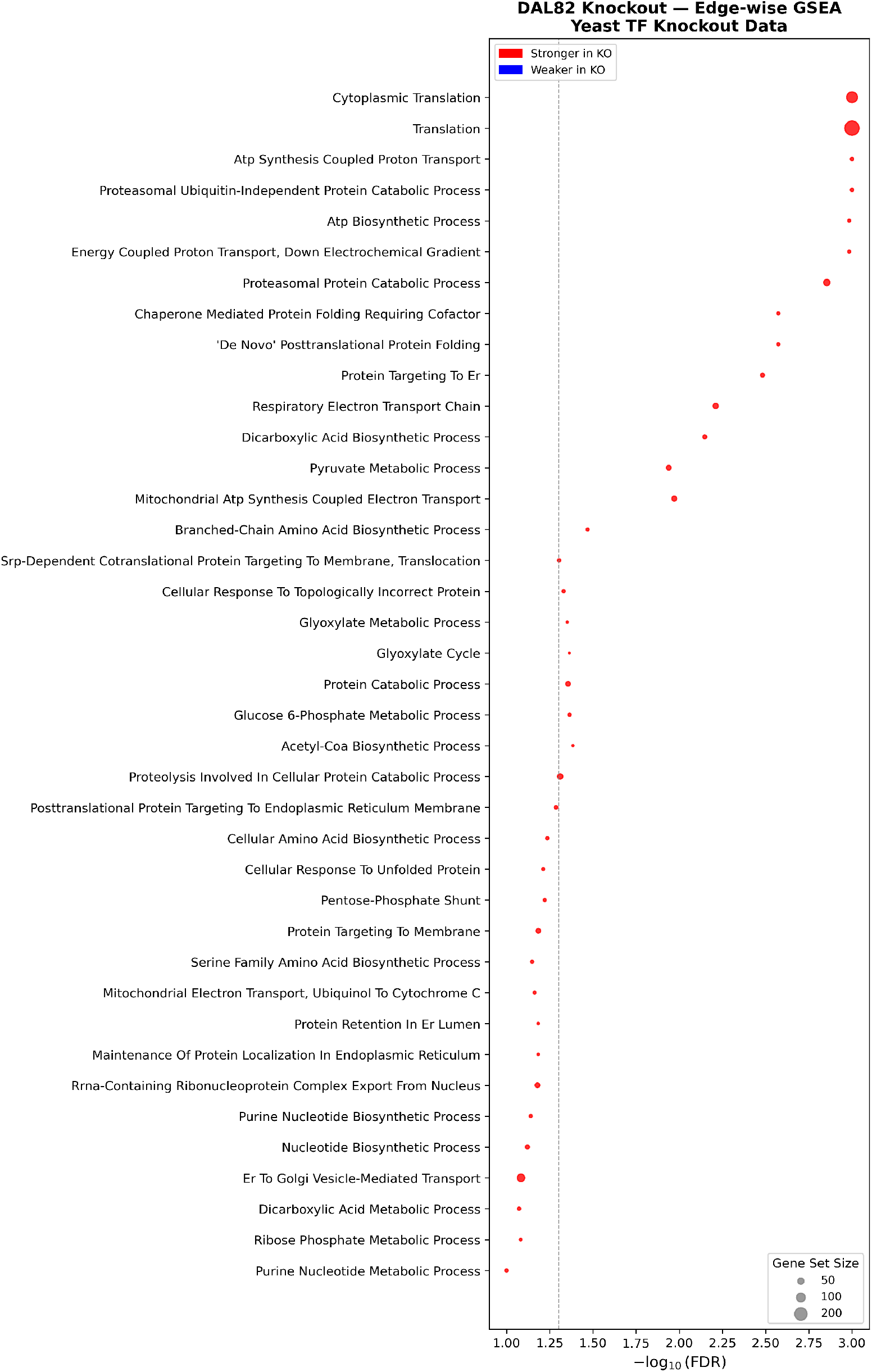
Gene set enrichment analysis of edges differentially connected to DAL82 in DAL82 knockout versus wild-type yeast samples. See Figure S2 for details.

**Figure S7.**
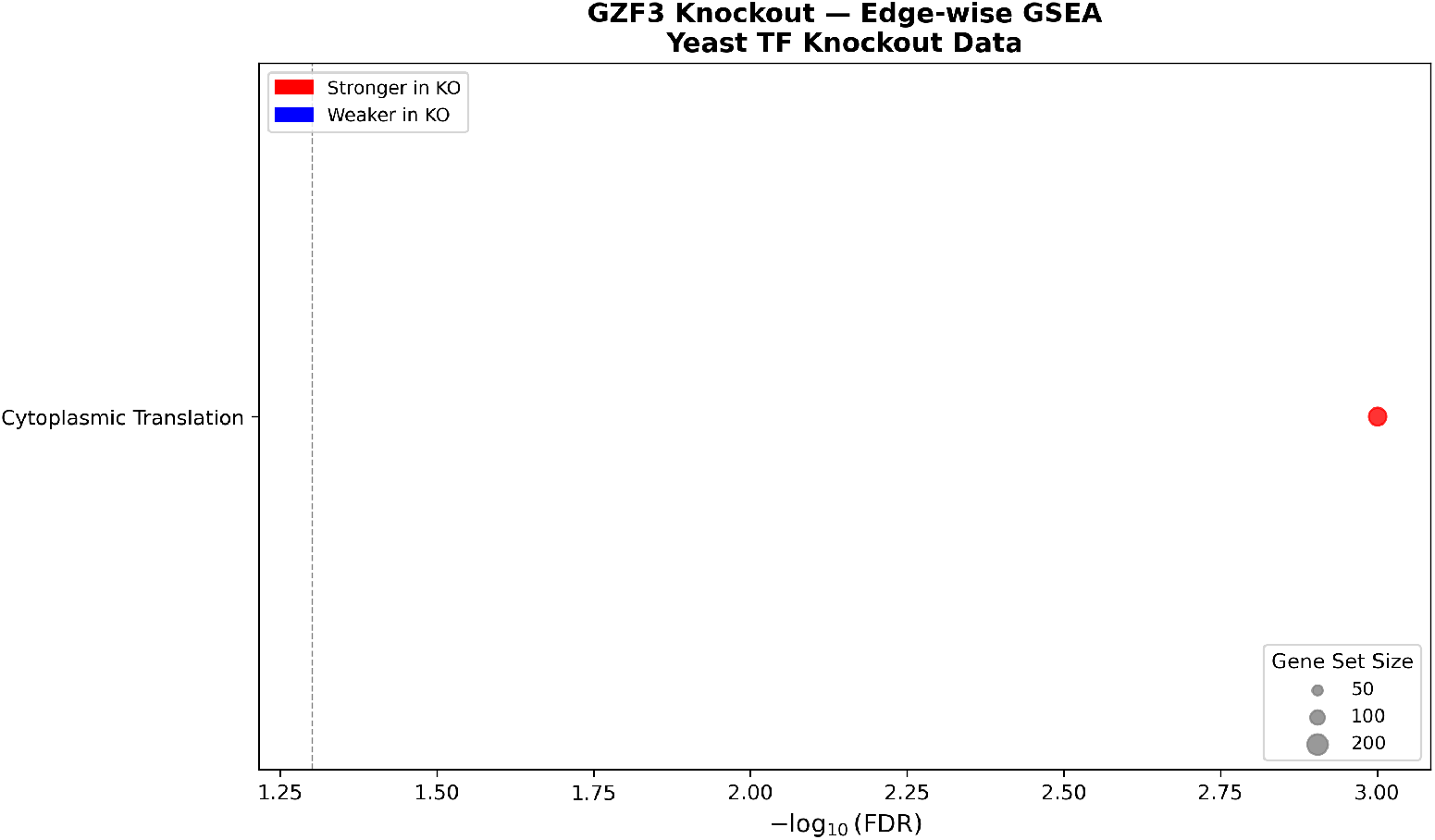
Gene set enrichment analysis of edges differentially connected to GZF3 in GZF3 knockout versus wild-type yeast samples. See Figure S2 for details.

**Figure S8.**
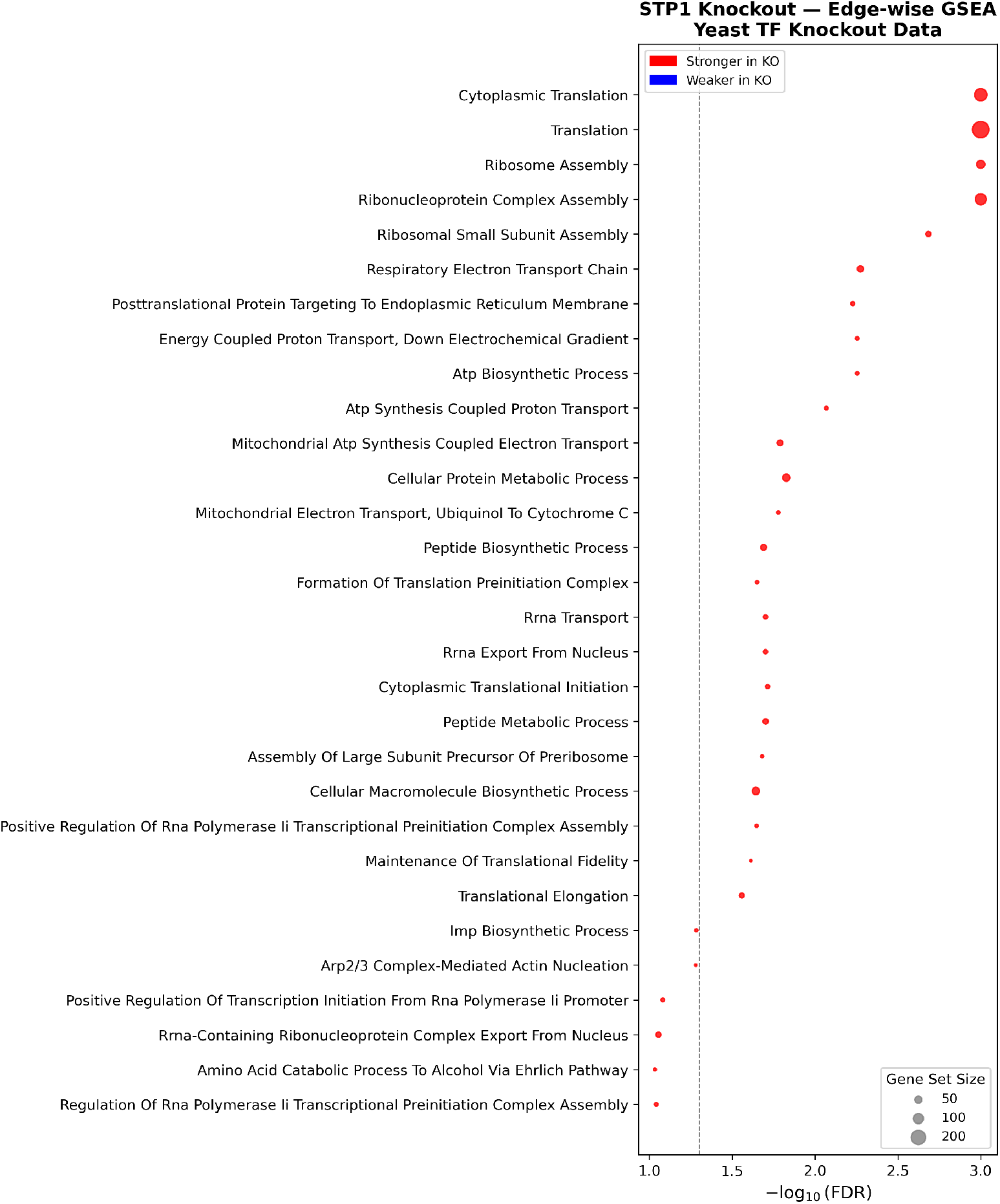
Gene set enrichment analysis of edges differentially connected to STP1 in STP1 knockout versus wild-type yeast samples. See Figure S2 for details.

**Figure S9.**
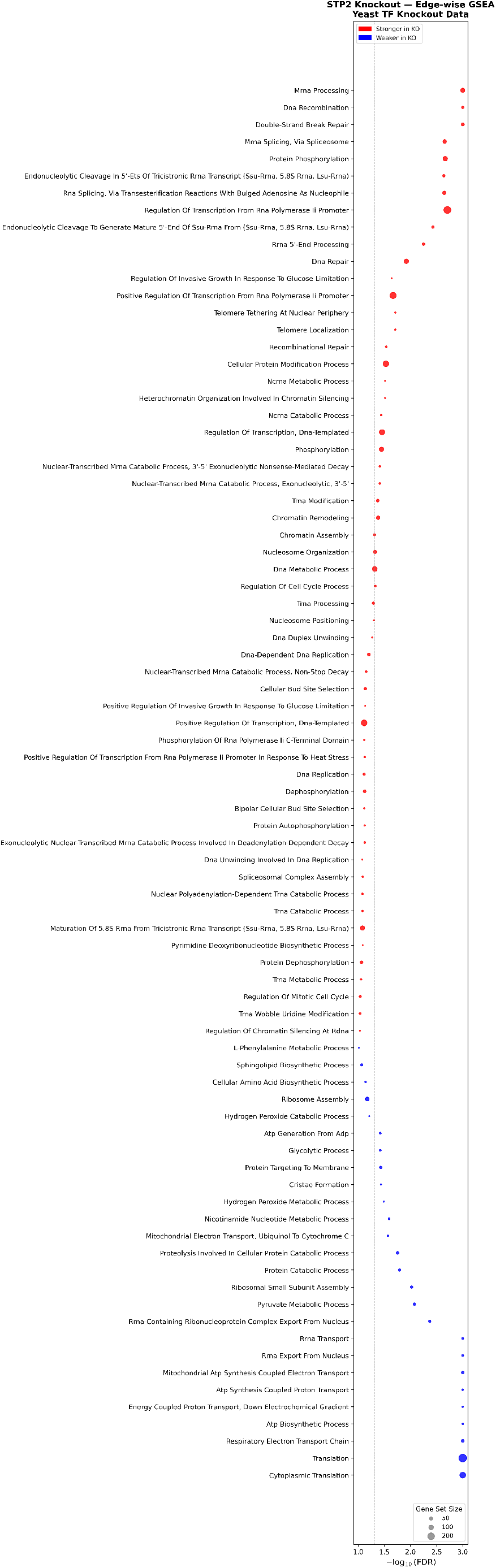
Gene set enrichment analysis of edges differentially connected to STP2 in STP2 knockout versus wild-type yeast samples. See Figure S2 for details.

**Figure S10.**
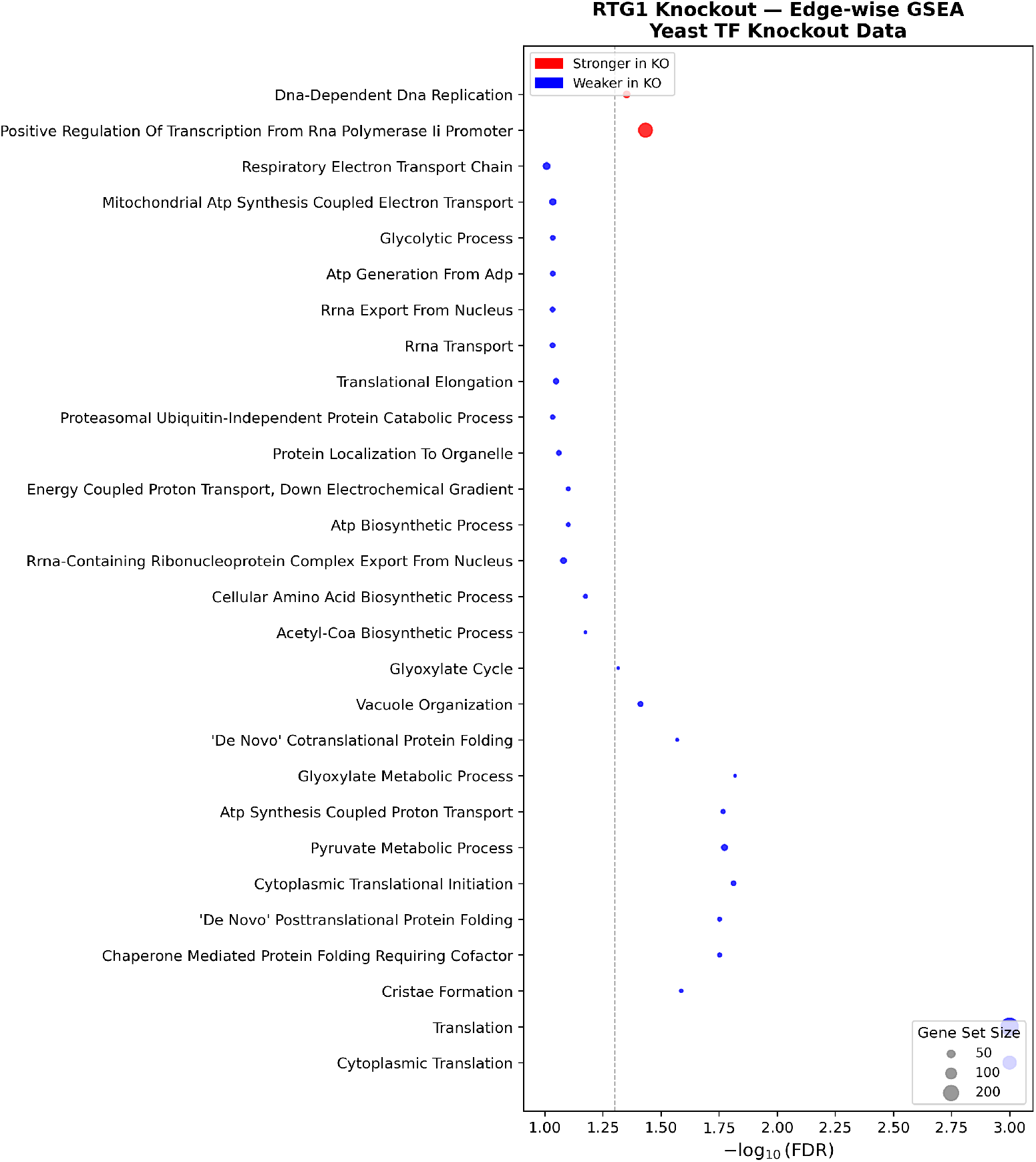
Gene set enrichment analysis of edges differentially connected to RTG1 in RTG1 knockout versus wild-type yeast samples. See Figure S2 for details.

**Figure S11.**
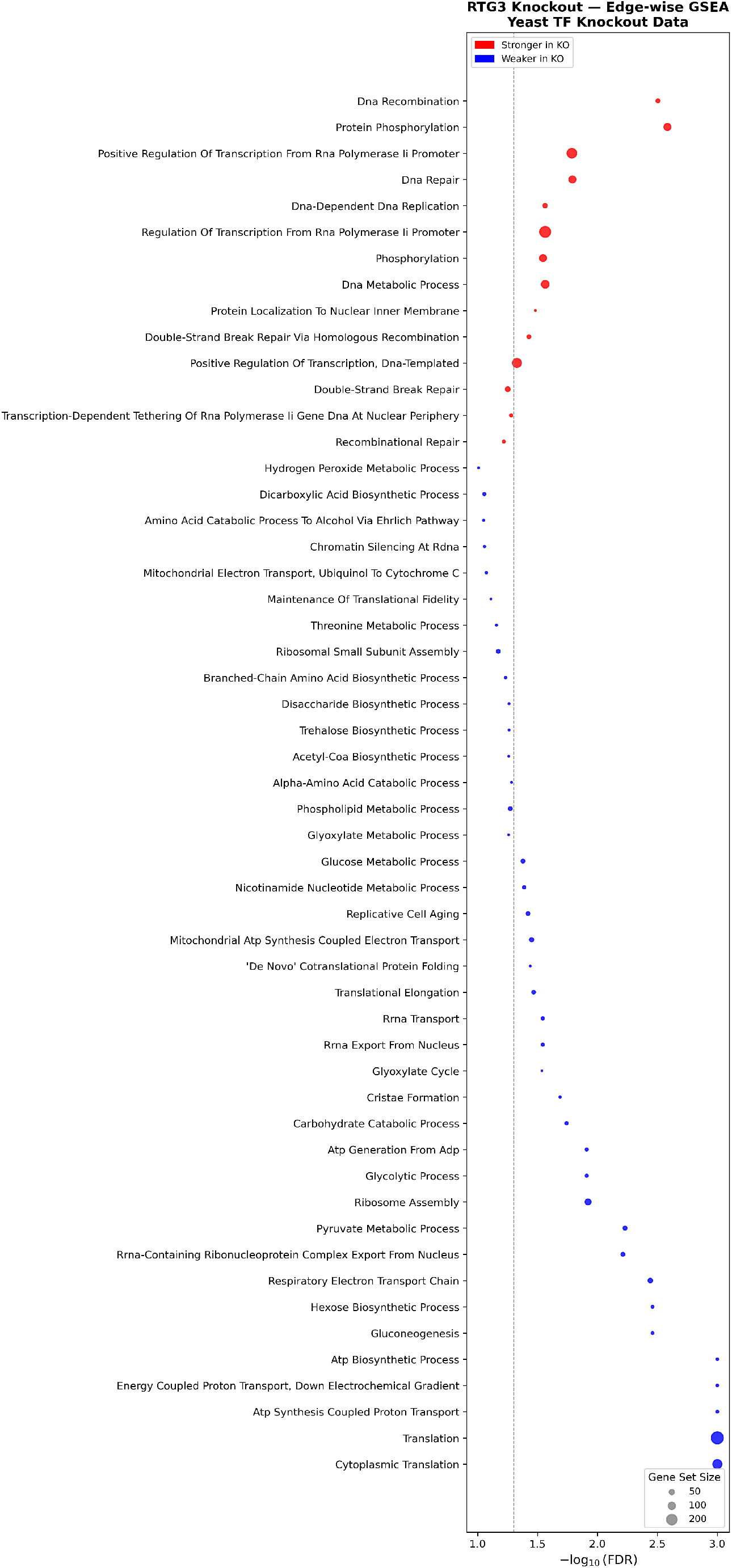
Gene set enrichment analysis of edges differentially connected to RTG3 in RTG3 knockout versus wild-type yeast samples. See Figure S2 for details.

### S4 Supplementary Figures for TCGA LUAD Data

**Figure S12.**
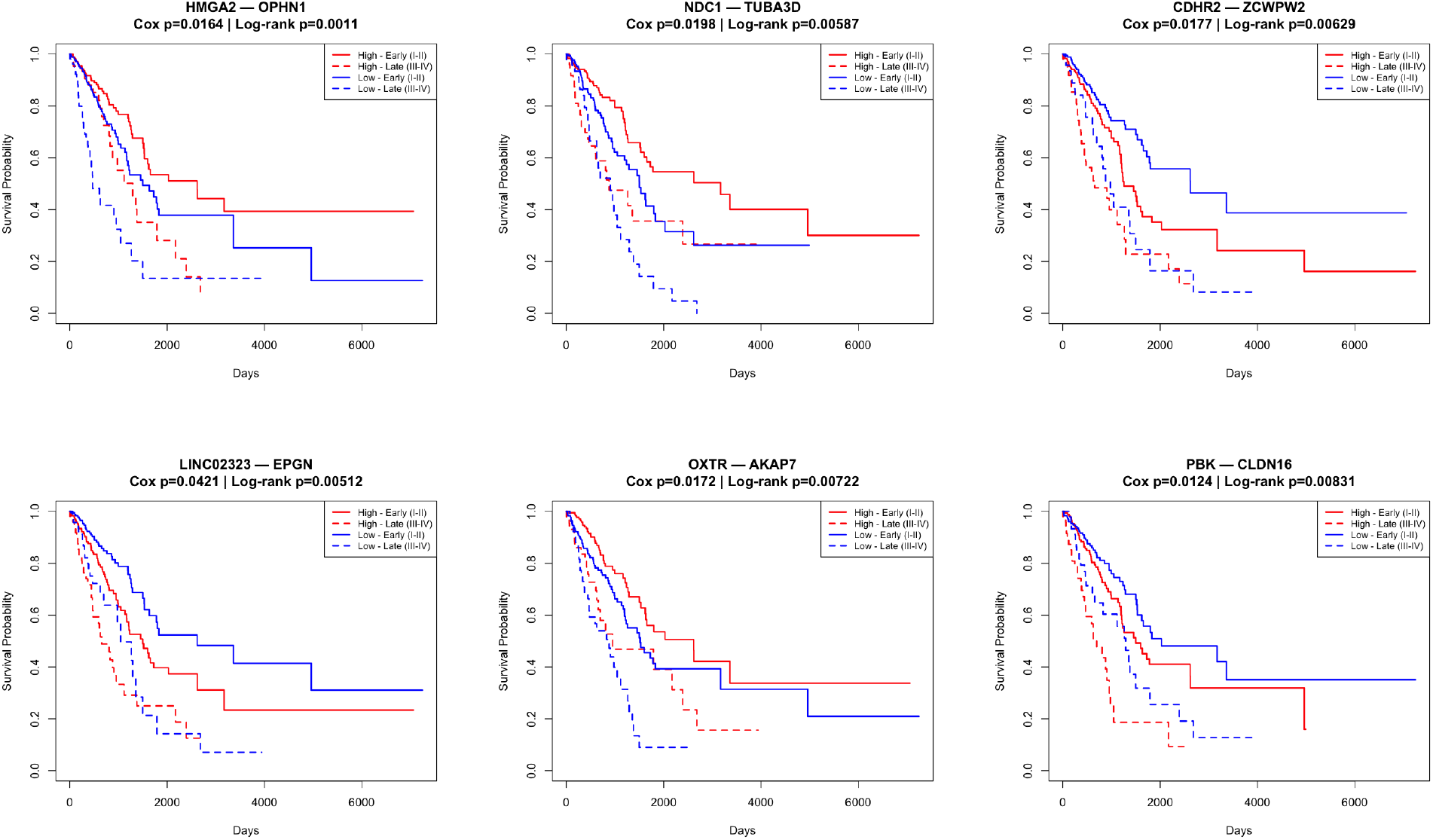
Kaplan-Meier survival curves for six additional representative survival-associated expression-methylation edges in lung adenocarcinoma. For each edge, patients are stratified by the median individual-specific partial correlation weight into high (red) and low (blue) groups, with solid and dashed lines representing early (Stage I-II) and late (Stage III-IV) disease respectively. Cox model p-values are adjusted for age, sex, tumor stage, smoking status, and race. Log-rank p-values are stratified by tumor stage.

